# Positive selection on a regulatory insertion-deletion polymorphism in *FADS2* influences apparent endogenous synthesis of arachidonic acid

**DOI:** 10.1101/042549

**Authors:** Kumar S.D. Kothapalli, Kaixiong Ye, Maithili S. Gadgil, Susan E. Carlson, Kimberly O. O’Brien, Ji Yao Zhang, Hui Gyu Park, Kinsley Ojukwu, James Zou, Stephanie S. Hyon, Kalpana S. Joshi, Alon Keinan, J. Thomas Brenna

## Abstract

Long chain polyunsaturated fatty acids (LCPUFA) are bioactive components of membrane phospholipids and serve as substrates for signaling molecules. LCPUFA can be obtained directly from animal foods or synthesized endogenously from 18 carbon precursors via the *FADS2* coded enzyme. Vegans rely almost exclusively on endogenous synthesis to generate LCPUFA and we hypothesized that an adaptive genetic polymorphism would confer advantage. The rs66698963 polymorphism, a 22 bp insertion-deletion within *FADS2*, is associated with basal *FADS1* expression, and coordinated induction of *FADS1* and *FADS2 in vitro*. Here we determined rs66698963 genotype frequencies from 234 individuals of a primarily vegetarian Indian population and 311 individuals from the U.S. A much higher I/I genotype frequency was found in Indians (68%) than in the U.S. (18%). Analysis using 1000 Genomes Project data confirmed our observation, revealing a global I/I genotype of 70% in South Asians, 53% in Africans, 29% in East Asians, and 17% in Europeans. Tests based on population divergence, site frequency spectrum and long-range haplotype consistently point to positive selection encompassing rs66698963 in South Asian, African and some East Asian populations. Basal plasma phospholipid arachidonic acid status was 8% greater in I/I compared to D/D individuals. The biochemical pathway product-precursor difference, arachidonic acid minus linoleic acid, was 31% and 13% greater for I/I and I/D compared to D/D, respectively. Our study is consistent with previous *in vitro* data suggesting that the insertion allele enhances n-6 LCPUFA synthesis and may confer an adaptive advantage in South Asians because of the traditional plant-based diet practice.

## Introduction

Twenty and twenty two carbon long chain polyunsaturated fatty acids (LCPUFA), especially arachidonic acid (ARA; 20:4n-6), eicosapentaenoic acid (EPA; 20:5n-3) and docosahexaenoic acid (DHA; 22:6n-3) are ubiquitous in mammalian tissue, are bioactive components of membrane phospholipids (PL) and serve as precursors to cell signaling eicosanoids and docosanoids that are major drug targets (e.g., COX-1, COX-2 inhibitors, leukotriene receptor antagonists) (Park, Kothapalli, Lawrence, et al. 2009; Park, Kothapalli, Reardon, et al. 2009; Park, et al. 2011). LCPUFA can be obtained directly from animal foods or endogenously synthesized from 18 carbon essential fatty acid precursors linoleic acid (LA; 18:2n-6) and alpha-linolenic acid (ALA; 18:3n-3) and their metabolites by an alternating series of desaturation and elongation reactions (Park, Kothapalli, Lawrence, et al. 2009). Vegans rely on this biochemical pathway to generate all LCPUFA from precursors. Classic carnivores (e.g. cats and most marine fish) have lost the metabolic ability to make LCPUFA and rely on consumption of animal tissue to supply all their LCPUFA requirements. Traditional human populations have been described that are analogous to herbivores (vegans) and carnivores (Natives of the Canadian Arctic) (Gibson and Sinclair 1981). The genomic determinants of the function of this pathway are therefore candidates for selective pressure based on human diets habitually consumed through many generations, e.g., by traditional vegan/vegetarian populations as are found in India. Signals of positive selection surrounding the FADS gene cluster have been reported in Greenlandic Inuit and in African populations (Ameur, et al. 2012; Mathias, et al. 2012; Fumagalli, et al. 2015).

The fatty acid desaturase genes *(FADS1* [OMIM#606148] and *FADS2* [OMIM#606149]) code for enzymes that catalyze the introduction of double bonds at specific positions in a fatty acid chain. FADS1 (Δ5-desaturase) and FADS2 (Δ6/Δ8/Δ4-desaturase) have specificity for several fatty acid substrates (Park, Kothapalli, Lawrence, et al. 2009; Park, et al. 2011; Park, et al. 2015; Park, et al. 2016). In humans, the FADS genes *(FADS1, FADS2*, and *FADS3)* evolved evolutionarily by gene duplication events and are clustered within the 100 kb region on the long arm of human chromosome 11 (11q12-13.1) (Marquardt, et al. 2000), a genomic cancer hot spot (Park, et al. 2011). Our previous results show extensive splicing of all three desaturases, including a novel function of an alternatively spliced isoform (Park, Kothapalli, Reardon, et al. 2009; Park, et al. 2010; Park, et al. 2012; Park, et al. 2015).

*FADS2* codes for a desaturase [EC 1.14.19.3] catalyzing the rate limiting steps in the biosynthesis of LCPUFA (Park, Kothapalli, Lawrence, et al. 2009). Supplementary fig. S1 shows the n-3 and n-6 biochemical LCPUFA pathways. Genetic studies have shown common single nucleotide polymorphisms (SNPs) within FADS gene cluster are strongly associated with LCPUFA levels and disease phenotypes (Schaeffer, et al. 2006; Malerba, et al. 2008; Tanaka, et al. 2009; Illig, et al. 2010). Using lymphoblasts from Japanese HapMap participants, we identified a 10 SNP-haplotype in *FADS2* (rs2727270 to rs2851682) controlling basal *FADS1* mRNA expression; minor allele carriers showed lower basal *FADS1* mRNA expression in cultured lymphoblasts (Reardon, et al. 2012). A conserved genomic region within the haplotype contained predicted binding sites for a sterol regulatory element binding protein (SREBP). By amplifying a 629 bp portion flanking the sterol response element (SRE), we identified a polymorphic 22 bp insertion/deletion (indel) genetic variant (rs66698963), of which the deletion is the minor allele. Minor allele homozygotes (D/D) had significantly lower expression of *FADS1* than the I/I major allele homozygotes (Reardon, et al. 2012). Arachidonic acid is the immediate product of FADS1, leading directly to the hypothesis that individuals carrying D/D genotype have lower metabolic capacity to produce LCPUFA from precursors than I/I individuals.

Here we present the first experimentally determined allele and genotype frequencies for the indel rs66698963 using a primarily vegetarian population from Pune, India compared to a U.S. population drawn broadly from around the country. By using genetic variations data from the 1000 Genomes Project (1000GP), we estimated the global genotype frequency distribution of rs66698963. By applying a series of tests for recent positive selection based on population divergence, site frequency spectrum (SFS), and long-range haplotype, we provide strong evidence that positive selection drove the insertion allele to high frequency not only in Africans, but also in South and some East Asians. The association of basal arachidonic acid status and indel genotype was evaluated in a subset of the U.S. population.

## Results and Discussion

### Experimentally Determined Genotype and Allele Frequencies of rs66698963

In the U.S. sampling, D/D genotype was 43% (n=134), I/D 39% (n=120) and I/I 18% (n=57); in the Indian samples D/D genotype was 3% (n=7), I/D 29.5% (n=69) and I/I 67.5% (n=158) (fig. 1A). The observed genotype frequencies in the U.S. sampling deviated from the Hardy Weinberg Equilibrium (HWE) expected frequencies, whereas sampling from India were consistent with HWE. As the U.S. population includes many definable subgroups, the deviation is likely to be due to stratification (Nussbaum, et al. 2007). In U.S. sampling, the observed allele frequency was D: 0.62, I: 0.38, whereas in Indian samples it was D: 0.18, I: 0.82. As a large fraction of our US sampling (n=201) is from the Kansas cohort, we tested allele frequency separately from this cohort and found similar trends (D: 0.64, I: 0.36).

**Figure 1.**
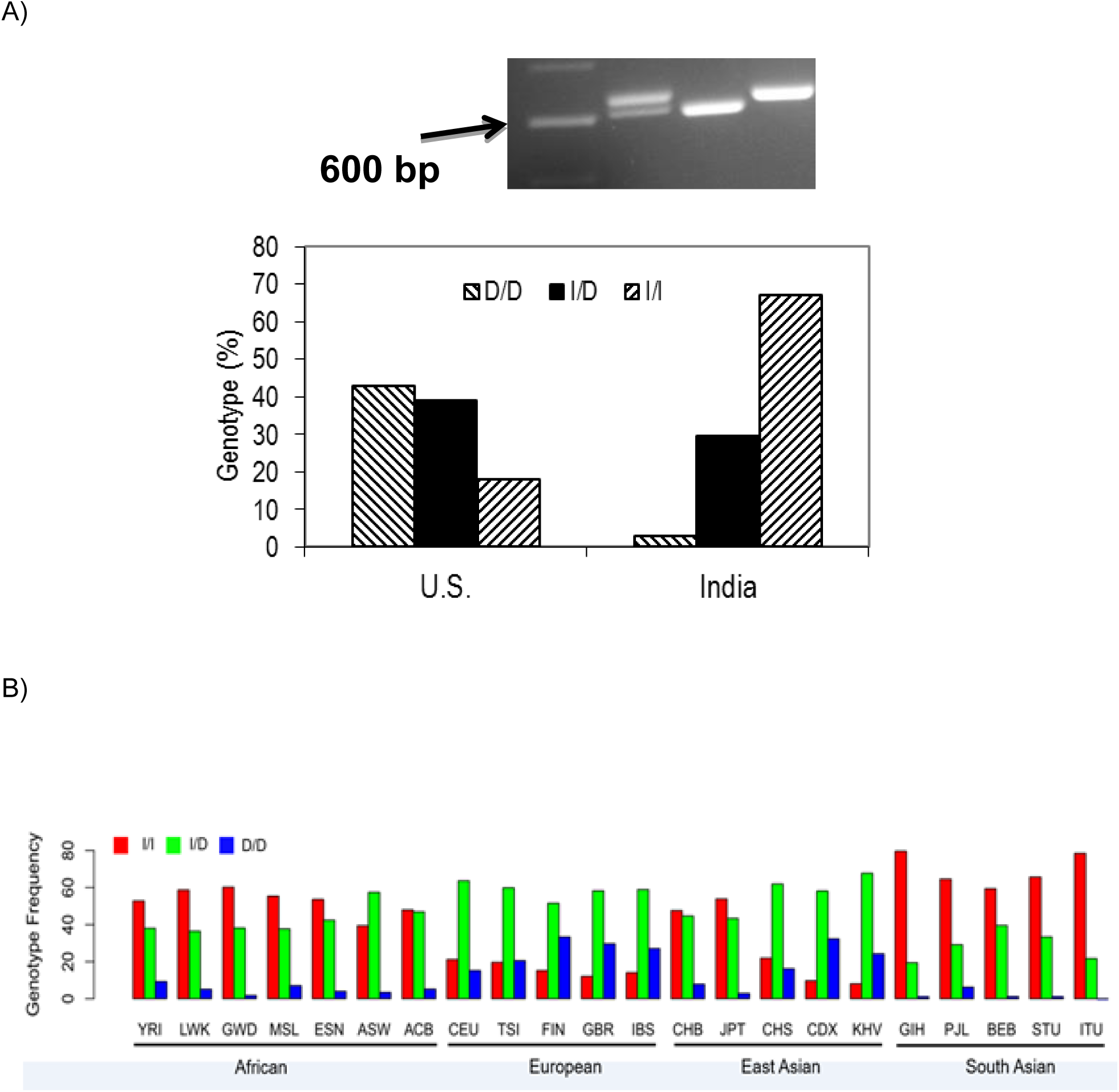
A) Top: Agarose gel image of PCR products from 3 individual DNA samples to establish genotype of the 22 bp rs66698963 polymorphism. Lanes from left to right: 100 bp ladder marker (arrow pointing at 600 bp), insertion/deletion (I/D); deletion/deletion (D/D); insertion/insertion (I/I). Botton: Distribution of rs66698963 allele frequencies in U.S and Indian subjects. B) Global genotype frequency distribution of rs66698963. Data was retrieved from the 1000 Genome Project (Materials and Methods).

### Global Genotype Frequency of rs66698963 Determined by using 1000GP Data

As our small sampling represents only two populations, we examined the global genotype frequency distribution of the rs66698963, by using the most recent whole-genome sequencing data from the 1000GP (Phase 3) (Auton, et al. 2015). The frequency of rs66698963 (indel) was retrieved from the 1000GP as the frequency of rs373263659 (indel) after we confirmed with Sanger sequencing that rs373263659 was actually a misannotation of rs66698963 (supplementary fig. S2). 1000GP data catalogs human genetic variation of 26 different global populations. Among the five populations of South Asian ancestry, D/D was found in 1.8% (n=9), I/D 28.2% (n=138), and I/I 69.9% (n=342), very consistent with observations from our experimental data from Pune, India. For the seven populations of African ancestry, D/D was 5.1% (n=34), I/D 41.5% (n=274), and I/I 53.4% (n=353). For the five populations of European ancestry, D/D was 25.0% (n=126), I/D 58.4% (n=294), and I/I 16.5% (n=83). For the five populations of East Asian ancestry, D/D was 16.3% (n=82), I/D 54.9% (n=277), and I/I 28.8% (n=145). Overall, the I/I genotype frequency is highest in African and South Asian populations, but lower in European and East Asian populations (supplementary table S1 and fig. 1B). Among the five populations of South Asian ancestry, Gujarati Indian from Houston, Texas (GIH) has the highest I/I genotype frequency, 80%, followed closely 78% by Indian Telugu from the UK (ITU). The lowest frequency of 59% is observed in Bengali from Bangladesh (BEB).

### Evaluation of Positive Selection on the FADS Region Surrounding rs66698963

Dramatic genotype frequency differences of rs66698963 among continental populations, led us to check if positive selection on the FADS region surrounding the rs66698963 has taken place especially in South Asian populations. We assessed rs66698963 genotype frequency differences among continental populations using the F_*ST*_ statistic, which quantifies the degree of population differentiation in allele frequencies. A value of 0 indicates no differentiation and value of 1 indicates complete subdivision. Over the four continental populations (African, European, East Asian and South Asian), the F_*ST*_ statistic of rs66698963 is 0.121 (empirical *p* value = 0.039), suggesting higher population differentiation than expected by chance. To gain insight into which population has undergone unusual frequency change as a result of regional adaptation, we further calculated pairwise F_*ST*_ among each pair of the four continental populations, as well as population branch statistics (PBS) for each population. Significantly high F_*ST*_ values were observed between South Asians and Europeans (F_*ST*_ = 0.276, empirical *p* = 2.71x10^−4^), between South Asians and East Asians (F_*ST*_ = 0.168, empirical p = 0.016), and between Africans and Europeans (F_*ST*_ = 0.157, empirical *p* = 0.044). No significant divergence was observed between South Asians and Africans (F_*ST*_ = 0.028) or between Europeans and East Asians (F_*ST*_ = 0.021). Together with the PBS patterns (supplementary fig. S3), these results suggest that rs66698963 has been subjected to positive selection in more than one of the studied populations.

To further test for positive selection signals in the region surrounding rs66698963 and to identify which populations have experienced positive selection, we applied formal tests of natural selection of two general types: (i) based on site frequency spectrum (SFS); and (ii) based on haplotype length and frequency. For SFS-based tests, we calculated genetic diversity as the average pairwise number of differences in the locus (π), Tajima’s D (Tajima 1989), and Fay and Wu’s H (Fay and Wu 2000). Positive selection could result in loss of genetic diversity, an excess of rare variants, which can be detected by Tajima’s D (negative values), and an excess of high-frequency derived alleles, which can be detected by Fay and Wu’s H (negative values). We applied these tests in each of the four continental regions, combining populations within the same continental region, and in each of the five populations of South Asian origin. All tests were performed across the whole genome using a sliding-window approach with a window size of 5 kilobase (Kb) and a step size of 1 Kb. Statistical significance was assessed with the empirical genome-wide distribution. Our analysis at the continental level (fig. 2A, supplementary figs. S4-S7) confirmed a pattern observed in a previous study: positive selection signals on *FADS1* was observed only in African populations, but not in European or East Asian populations (Mathias, et al. 2012). For this genomic region surrounding *FADS1*, the three test statistics in the South Asian populations follow a similar but less significant pattern as in African populations. For the immediate genomic region surrounding rs66698963, we observed extremely negative Fay and Wu’s H values, restricted to South Asians (fig. 2A). African populations also had lower H values in the same region, but the reduction was not significant. For the combined group of five populations of South Asian origin, three consecutive 5-Kb windows, representing chromosome 11 region 61,601,001 – 61,608,001 bp, had significant Fay and Wu’s H values, with the most significant value being −12.27 (empirical *p* = 0.031). For each of the five populations individually, we also observed significant H statistics surrounding rs66698963 (fig. 2B, supplementary figs. S8-S12). Two of these populations, GIH and ITU, exhibited the most significant signals of positive selection (supplementary figs. S9 and S10). The most significant H statistic in GIH was −14.20 (empirical *p* = 0.015) and in ITU it was −13.80 (empirical *p* = 0.018). The Tajima’s D statistic also showed a significant signal of positive selection for these two South Asian populations: −1.99 in GIH (empirical p = 0.013) and −1.90 in ITU (empirical *p* = 0.025).

**Figure 2.**
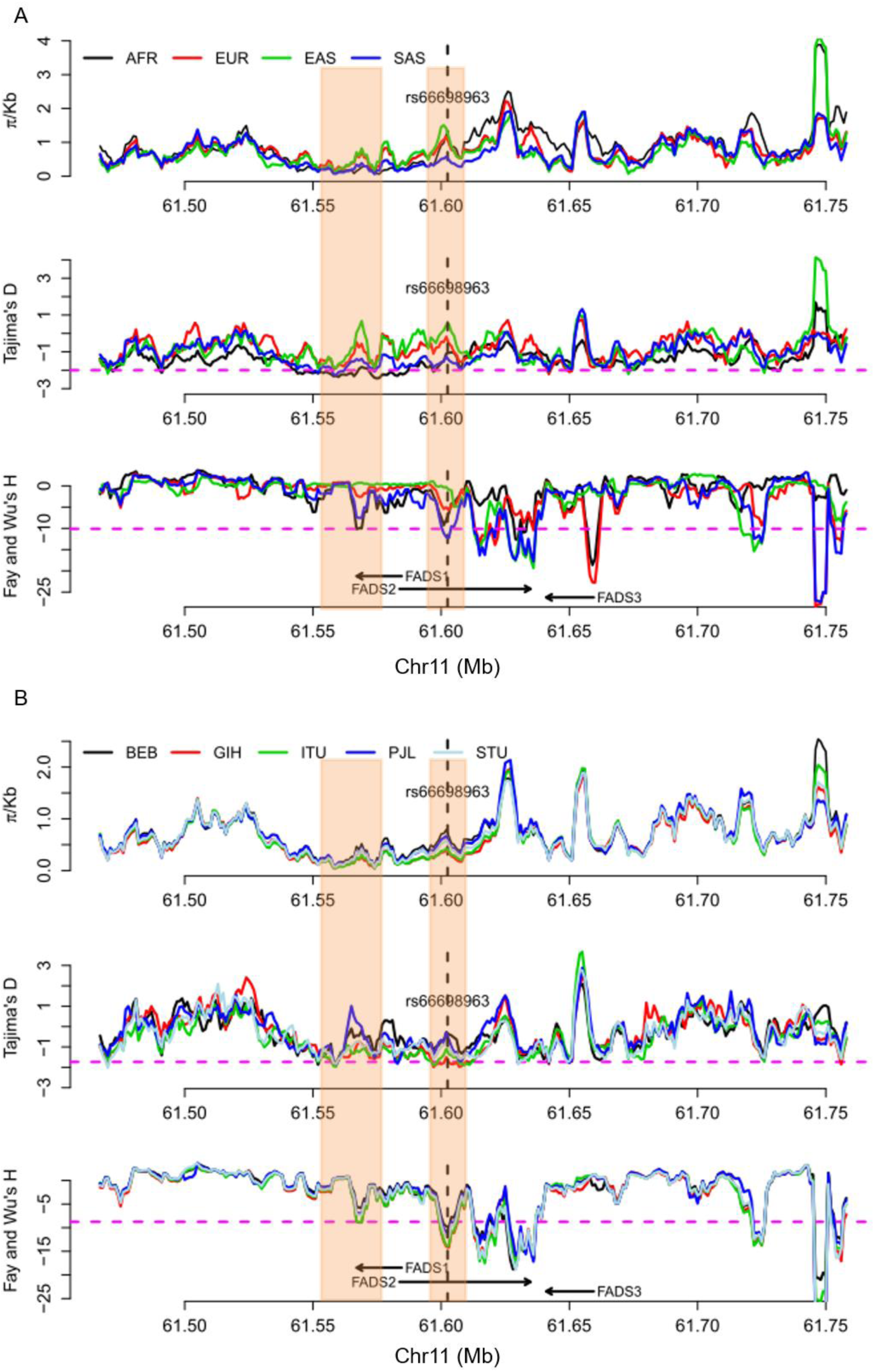
Patterns of genetic diversity (π), Tajima’s D, and Fay and Wu’s H across a 300 Kb genomic region around FADS genes in *A)* four continental regions, each combining multiple populations from the same regions; and *B)* five populations of South Asian origin. The three statistics were calculated using a sliding window method with window size of 5 Kb and moving step of 1 Kb. The left shaded box indicates the region that have been found (and confirmed in this study) to be under positive selection in African populations. The right shaded box indicates the genomic region that has significant SFS-based test results in South Asians, but not in others. The pink dashed lines in *A)* represent an empirical *p* value of 0.05 based on statistics from all genome-wide windows in the South Asian (SAS) continental region. Note that the statistical cutoffs vary slightly in different continental regions. The pink dashed lines in *B)* represent the lowest 5% cutoff among the five populations, that is, all genomic regions below the pink dashed line reach statistical significance regardless of the population. Note that some genomic regions above the pink dashed line could still be significant in some populations because the 5% cutoffs in these populations are higher (less extreme).

For haplotype-based tests for positive selection, we calculated the integrated Haplotype Score (iHS) (Voight, et al. 2006), the number of segregating sites by length (nSL) (Ferrer-Admetlla, et al. 2014), and Cross-population Extended Haplotype Homozygosity (XPEHH) (Sabeti, et al. 2007) for all genome-wide common variants (minor allele frequency > 5%) and statistical significance of each statistic was assessed with the empirical genome-wide distribution. Among the four continental regions (fig. 3), the most significant signals of positive selection were observed in South Asians, which have an iHS value of −2.79 (empirical *p* = 4.7x10^−3^) and a nSL value of −4.38 (empirical *p* = 8.95x10^−5^), with the haplotype carrying the insertion allele showing unusually long length and high frequency (fig. 4). Significant XPEHH of rs66698963 was also observed between South Asians and Europeans with a value of 3.47 (empirical *p* = 1.2x10^−3^, fig. 5A). Taken together, these haplotype-based test results are consistent with the SFS-based tests and point to a selective sweep on the *FADS* region in South Asians, with the adaptive allele being the insertion of rs66698963 or a variant in very strong linkage disequilibrium (LD) with it.

**Figure 3.**
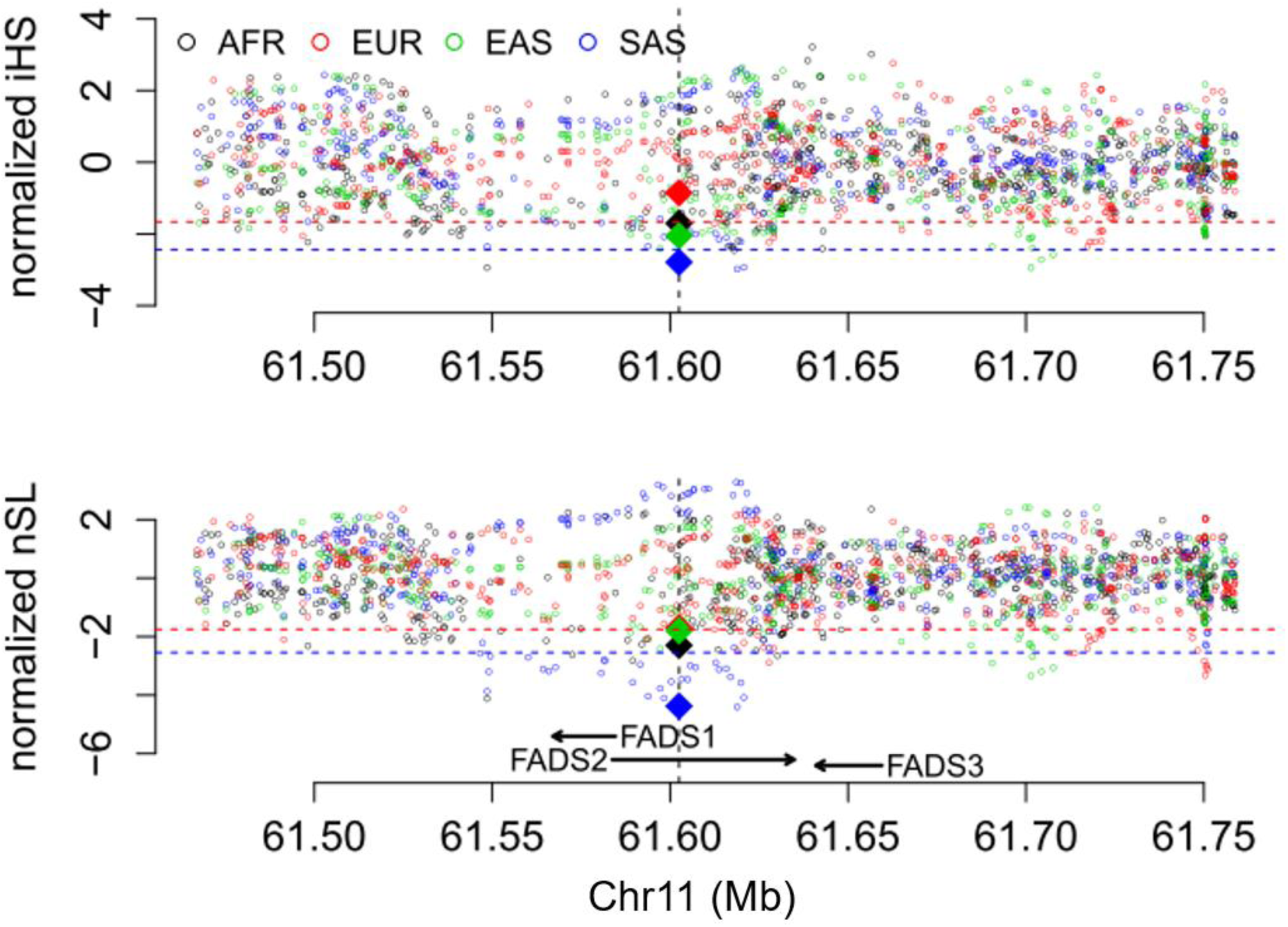
Patterns of normalized iHS and nSL (y-axes) across a 300 Kb genomic region (x-axis) around the FADS gene cluster in four continental regions, each combining multiple populations in the same region. The two statistics were calculated for each of the common SNPs (minor allele frequency > 5%) in the genomic region, as well as for the rs66698963 indel. The position of rs66698963 is indicated as a vertical black dashed line. The values for rs66698963 in each continental region are highlighted as diamonds filled with region-specific colors. The red dashed lines represent empirical *p* values of 0.05, while the blue dashed lines represent empirical *p* values of 0.01 based on estimates in the South Asian (SAS) continental region. Note that the statistical cutoffs vary slightly in different continental regions. Detailed representations for each continental region with their specific significant cutoffs are presented in supplementary figs. S13-S16.

**Figure 4.**
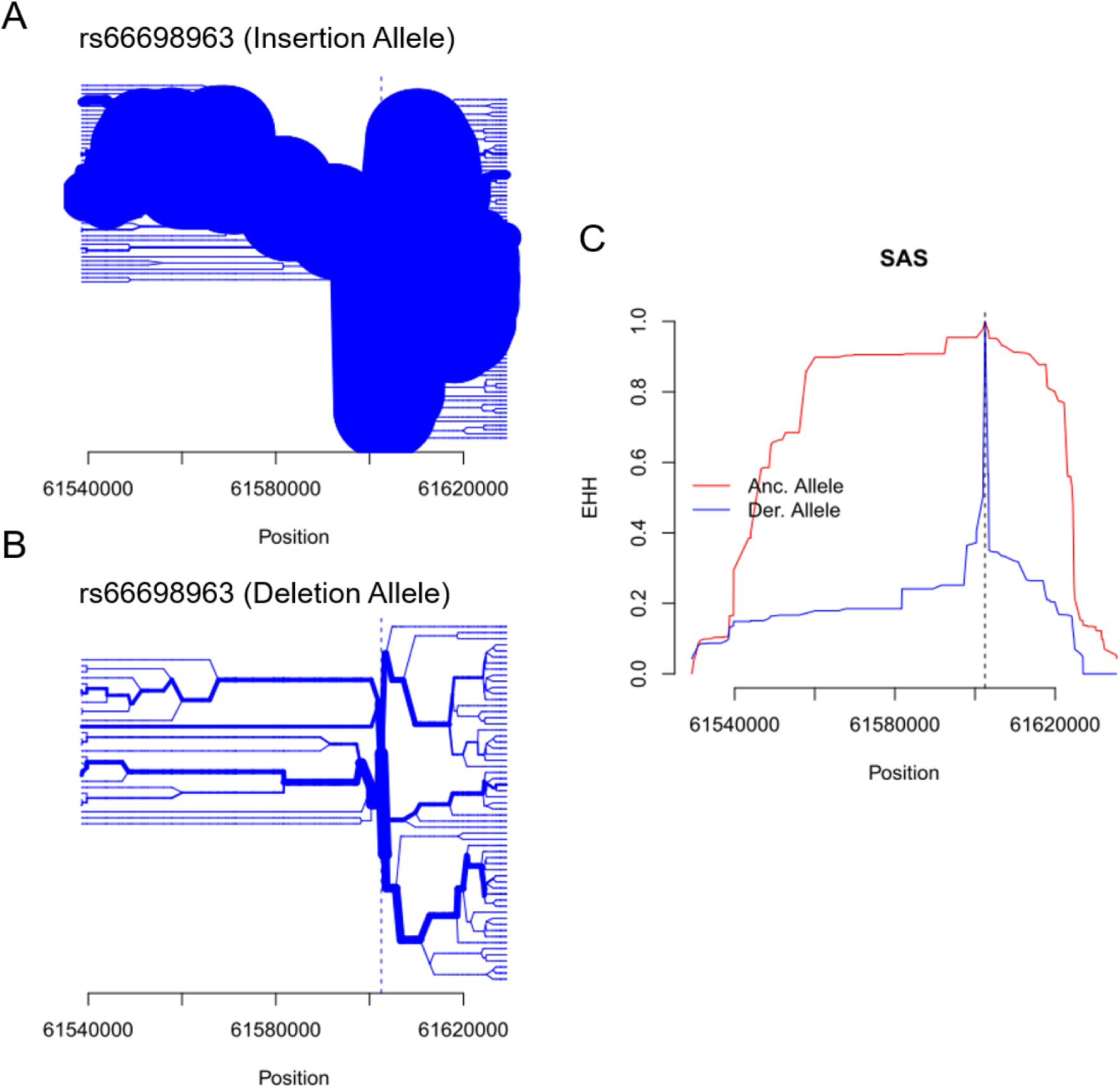
The insertion allele of rs66698963 is associated with signals of positive selection in the South Asian continental region. A) Haplotype bifurcation diagrams for the insertion allele at rs66698963; B) Haplotype bifurcation diagrams for the deletion allele at rs66698963; C) expanded haplotype homozygosity (EHH; y-axis) for rs66698963 (denoted by vertical dashed line). The EHH values are plotted against the physical distance extending both upstream and downstream of the target indel. The ancestral allele, also the insertion allele, shows much extended haplotype homozygosity than the derived allele, which is the deletion allele. Similar figures for AFR and EAS are presented in supplementary figs. S18 and S19.

In addition to South Asians, signals of positive selection were also found in Africans and East Asians (fig. 3, supplementary figs. S13-S19). For continental level analysis with populations in the same continental region combined into a single group, Africans have iHS value of −1.70 (empirical *p* = 0.043) and nSL value of −2.30 (empirical *p* = 0.017) for rs66698963 while East Asians have iHS value of −2.05 (empirical *p* = 0.022) and nSL value of −1.80 (empirical *p* = 0.045). No significant signals of positive selection were detected for Europeans in any of our tests. Consistent with population divergence results based on F_*ST*_, XPEHH of rs66698963 is also significant between Africans and Europeans with a value of 1.64 (empirical *p* = 0.039, fig. 5B).

**Figure 5.**
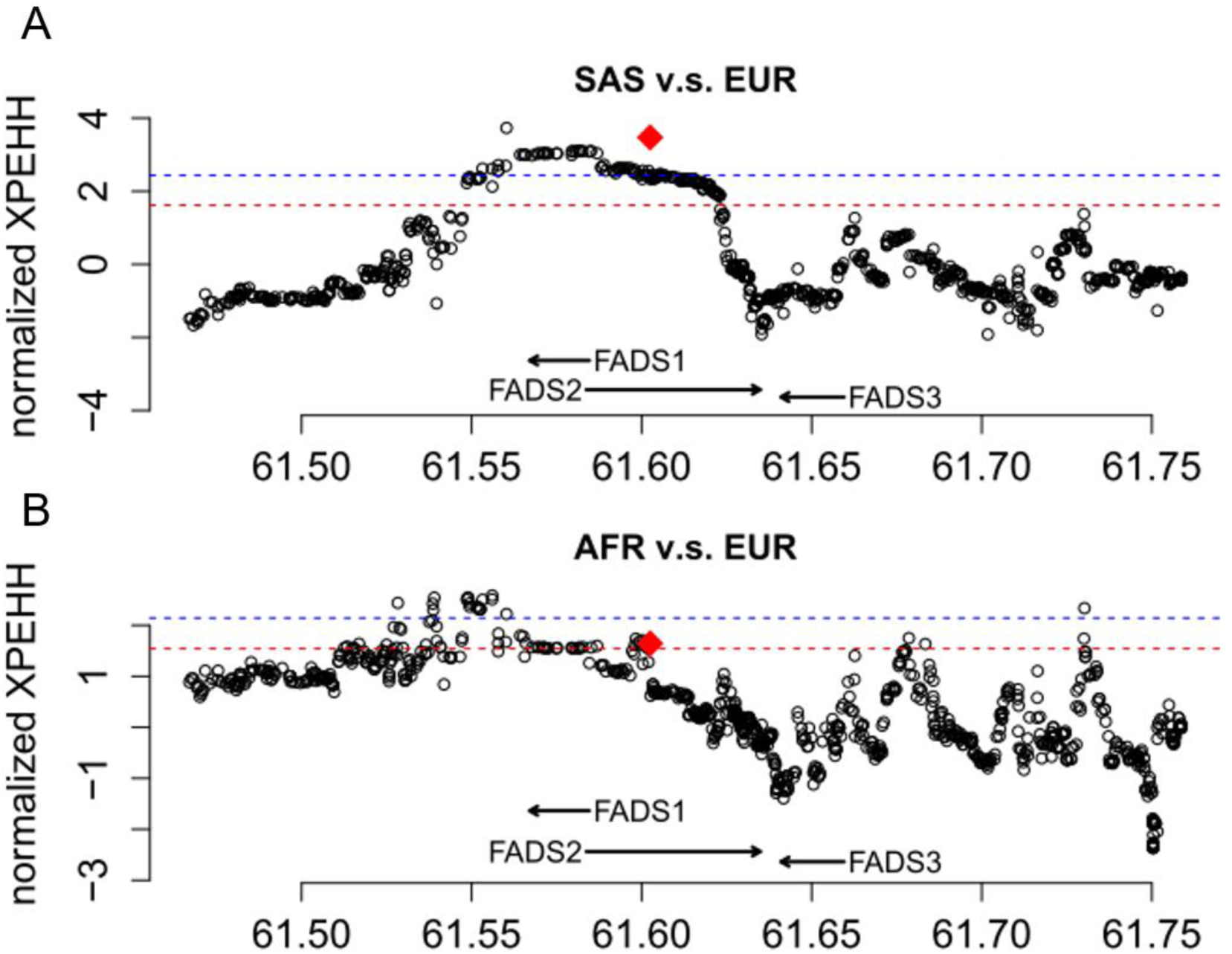
Patterns of normalized XPEHH (y-axes) across a 300 Kb genomic region around FADS genes A) between SAS and EUR, and B) between AFR and EUR. The normalized XPEHH statistic was calculated for each common SNP in the region, as well as for the rs66698963 indel. The value for rs66698963 is highlighted as a red diamond. The red dashed line and the blue dashed line represent empirical *p* values of 0.05 and 0.01, respectively.

Finally, we performed haplotype-based analyses in each of the five East Asian populations separately because of the frequency difference of rs66698963 across them (fig. 1B) and because analysis of East Asians combined as a single group produced apparent inconsistent results: significant haplotype-based results but lack of significant divergence from Europeans. Significant iHS and nSL results of rs66698963 were only found in CHB and JPT (supplementary fig. S17), the two populations with highest insertion allele frequency (70% and 75% respectively). Taken together, strong signals of positive selection on the region surrounding rs66698963 were consistently found in South Asians, Africans and some East Asians. All results point to rs66698963 as being at the center of the signal of selection, with the adaptive allele being the insertion allele of rs66698963 or a variant in very strong LD with it.

### Challenge in Indel Calling with Low-depth Sequencing Data

While our Sanger sequencing and agarose gel electrophoresis data provided accurate frequency estimates for the Indian and U.S. samples, the global frequency pattern of rs66698963 derived from the 1000GP data may be subjected to inaccurate annotation. Indel calling has been proved challenging with short-read mapping data sets and indel variants cataloging is still in its infancy (Li, et al. 2014). Due to the low sequencing coverage depth (~6 X) in the 1000GP, annotation error is possible. Using the Sanger sequencing data of rs66698963 from 16 Japanese (JPT) individuals from our previous study, which were chosen to be homozygotes at rs66698963 (Reardon, et al. 2012), we assessed the accuracy of frequency estimates from 1000GP. Based on Sanger sequencing, 11 out of 16 individuals are I/I genotype and the remaining five are D/D genotype. All eleven I/I individuals were accurately identified in 1000GP, whereas, among the five D/D individuals, only one was correctly identified, with the other four inaccurately annotated as I/D heterozygotes. This observation suggests that inaccurate allele calling in 1000GP may result in lower number of D/D individuals and higher number of I/D individuals (fig. 1B). Better sequencing data are needed in the future for accurate estimates. It is noteworthy that our above analysis of natural selection based on SFS and long-range haplotype are unlikely to be affected by the inaccuracy of rs66698963 calling, because these statistical tests draw information from many genetic variants in the surrounding region.

### Interpretation of the Signals of Positive Selection Surrounding rs66698963

Our evolutionary tests based on population divergence, SFS, and long-range haplotype, demonstrated consistently strong signals of positive selection surrounding rs66698963 in South Asian, African and some East Asian populations (CHB and JPT). Positive selection on the FADS gene cluster have been reported in Africans (Ameur, et al. 2012; Mathias, et al. 2012) and in Greenlandic Inuit (Fumagalli, et al. 2015) populations. Our study is the first to demonstrate positive selection in South and some East Asian populations. Most importantly, we propose a putatively causal indel, with regulatory effects on *FADS1* and *FADS2* (Reardon, et al. 2012).

The signatures of positive selection in African populations previously described by Mathias et al. (Mathias, et al. 2012) were replicated in our study and further confirmed with additional evidence from haplotype-based tests (fig. 3). This region is upstream of rs66698963 and its positive selection signatures are unlinked from those immediately surrounding the indel. However, the possibility could not be ruled out that the same adaptive variant is responsible for the observed signatures of selection in both clusters. The other study suggesting positive selection on FADS genes in African populations proposed an adaptive haplotype based on 28 SNPs in the FADS region (Ameur, et al. 2012). Although no positive selection signals were found in European populations, the adaptive haplotype in Africans, referred to as haplotype D, is present in 62% of Europeans and associated with greater LCPUFA status in Europeans (Ameur, et al. 2012). To test the possibility that the adaptive and association signals of this haplotype D are the same as those of the insertion allele of rs66698963, we examined the proportion of haplotype D carrying insertion allele at rs66698963. Among all phased haplotypes from the four continental regions, 2829 carry insertion allele at rs66698963, 72.9% of them also carry the haplotype D. For 2357 haplotypes carrying haplotype D, 87.5% also carry the insertion allele (supplementary fig. S20A). To mitigate the effect of annotation error at rs66698963, we further tested the correspondence between the haplotype D defined with 28 SNPs and the major haplotype defined with 10 SNPs from our previous study (Reardon, et al. 2012). These two sets of SNPs have three in common. Based on our previous study in Japanese individuals, all 10-SNP major haplotype carry insertion allele at rs66698963 (Reardon, et al. 2012). Among 2762 chromosomes carrying the major haplotype, 85.05% also carry haplotype D, while 99.66% of 2357 chromosomes carrying haplotype D also carry the major haplotype (supplementary fig. S20B). This strong correspondence between haplotype D and our major haplotype was also observed when stratifying into each of the four continental regions (supplementary fig. S20C-F), indicating that the association and adaptive signals described in Ameur et al. (Ameur, et al. 2012) are very likely the same as we reported in this study. However, since previous evidence of positive selection was only drawn from XPEHH and a composite likelihood (CLR) test of the allele frequency spectrum (Ameur, et al. 2012), our analyses, with a series of SFS-based and haplotype-based tests, dramatically strengthen the evidence of positive selection in Africans. Most importantly, our study proposes a putatively causal variant and demonstrates that the adaptive signals are also present in South and some East Asians, with the strongest signals in South Asians.

Most recently, Fumagalli et al. described signatures of adaptation on the FADS gene cluster in indigenous people of Greenland, the Inuit, which have subsisted on a marine diet rich in omega-3 PUFA (Fumagalli, et al. 2015). Two SNPs carrying adaptive signals, rs7115739 and rs174570, were also associated with reduced LCPUFA status. The derived alleles of these two SNPs are almost fixed in the Greenlandic Inuit but have much lower frequencies in other populations (Fumagalli, et al. 2015). We observed that one of the two SNPs, rs174570, also exhibits positive selection signals in South Asians (nSL = −3.77, p = 0.00048) and East Asians (iHS = −2.41, p = 0.010; nSL = −2.41, p = 0.012). Interestingly, it is the ancestral allele of the SNP that is adaptive in these populations, in contrast to the derived allele in Greenlandic Inuit (Fumagalli, et al. 2015). The ancestral allele has strong LD with the insertion allele of rs66698963 in South and East Asians (D’ = 0.72 and 0.92, respectively). It is plausible that the ancestral allele of rs174570 may be tagging the effect of the insertion allele of rs66698963. If that is the case, our study might not only report a functional, adaptive allele (the insertion) in South Asians, some East Asians, and Africans, traditionally residing on plant-based diet, but also an adaptive allele (the deletion) in Greenlandic Inuit with marine diet rich in omega-3 fatty acids.

### Functional Analysis of rs66698963 Genotypes and Fatty Acid Levels

Associations between genotype and PUFA concentration in RBC PL from the Kansas cohort were used to probe whether the I/I genotype, with putative higher endogenous arachidonic acid synthetic capacity, follow a pattern of metabolites consistent with the pathway: higher product and lower precursor in the order I/I, I/D, and D/D. RBC fatty acid levels were measured in 199 Kansas cohort subjects, of them n=89 were D/D genotype, n=76 were I/D genotype and n=34 were I/I genotype. Participant characteristics of the Kansas cohort are presented in supplementary table S2. n-6 PUFA acids were significantly and consistently related to genotype. Fig. 6 presents associations between genotype and n-6 PUFA concentration in RBC PL, a long term (several month) measure of whole body status. Linoleic acid (18:2) was not related to genotype. In contrast, the LCPUFA 20:3n-6, 20:4n-6 (arachidonic acid) and 22:4n-6 (adrenic acid) are all strongly related to genotype. All LCPUFA comparisons are significant between the D/D and I/I genotypes. 20:3n-6 levels are highest in D/D and lowest in I/I, whereas the 20:4n-6 and 22:4n-6 are opposite. The differences in PUFA concentration calculated for various precursor-product pairs shows the strongest relationship between 20:4n-6 and 18:2n-6, where I/D and I/I are both different from D/D. Other differences are not as strong but consistent with expectations from the desaturation-elongation pathway. All differences between genotypes remained significant after controlling for the effect of age.

**Figure 6.**
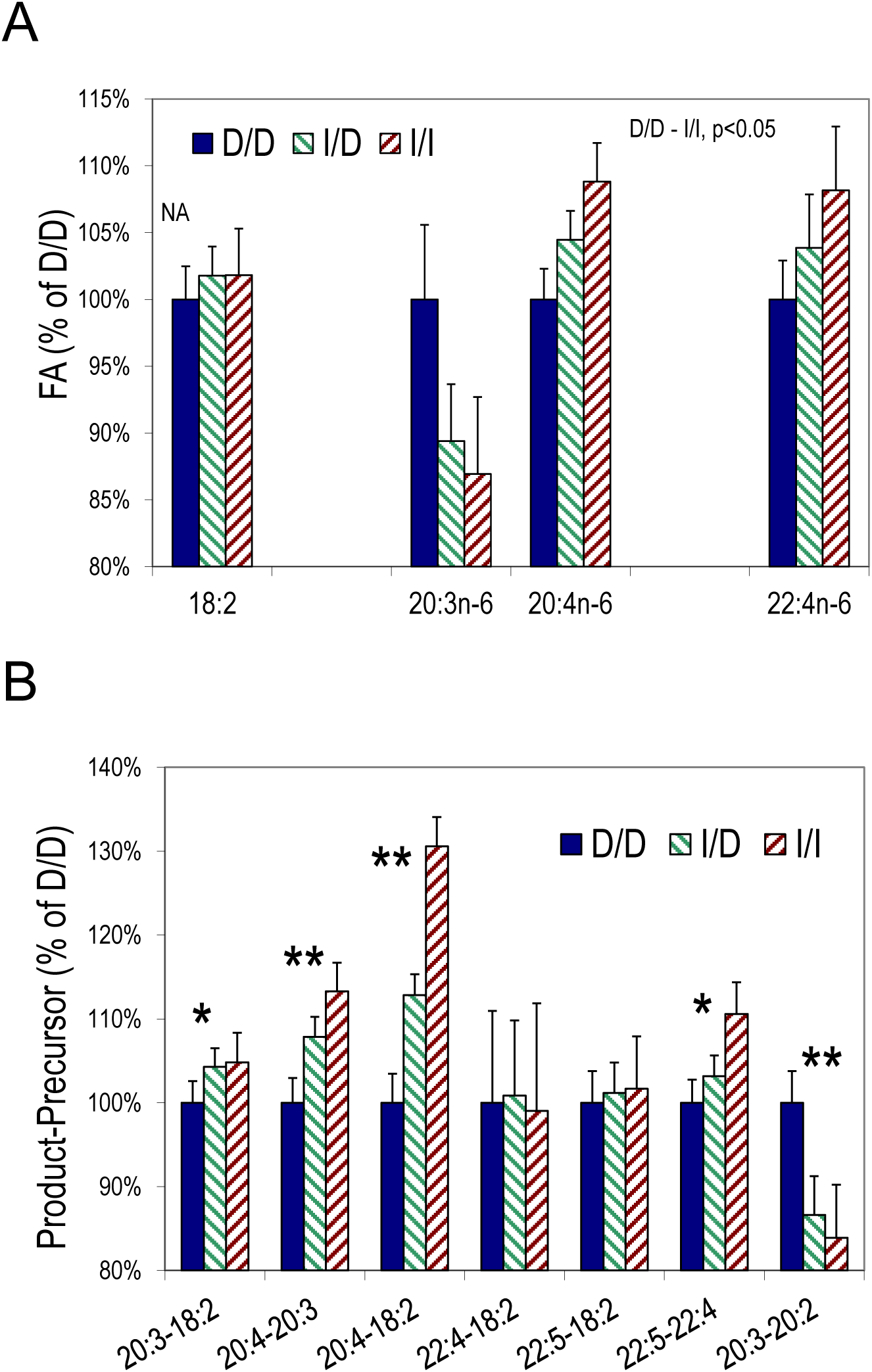
RBC n-6 PUFA as a function of genotype. PUFA in the biochemical pathway 18:2 (*FADS2+Elongation*) → 20:3 (*FADS1*) 20:4 → 22:4 (*FADS2*) → 22:5 where *FADS2* and *FADS1* proteins mediate the indicated transformations. Percent changes relative to the D/D genotype are calculated as A_x/x_/A_D/D_, where A is a 18:2, 20:3, 20:4, 22:4, and the units are %,w/w. *A)* 18:2 (linoleic acid) consumed exclusively by diet is not related to genotype. 20:3n-6 is lower but 20:4n-6 higher, consistent with a direct influence on 20:3→20:4 by *FADS1*. All comparisons are significant, except 18:2 (NA). *B)* Differences in concentration between upstream-downstream PUFA normalized to the D/D values. All genotype-dependent differences involving *FADS1* or *FADS2* mediated step(s). Percent differences calculated as [(B-A)_x/x_)]/[(B-A)_D/D_], where B is a downstream product and A is the precursor, and the units are %,w/w. By t-test, *I/I vs D/D p<0.05; ** I/I or I/D vs D/D p<0.05. By the non-parametric Wilcoxon method, all comparisons correspond to t-test results except: (A) 22:4 I/I vs D/D, p=0.09; (B) 20:3-18:2, p=0.07; 22:5-22:4, p=0.11; 20:3-20:2, p<0.05 for all pairwise comparisons.

The metabolic pathway relating these PUFA is 18:2n-6→→20:3n-6→20:4n-6→22:4n-6. The step 20:3→20:4 is mediated by *FADS1*, basal expression of which our previous data show is modulated by the genotype (Reardon, et al. 2012). The three steps between the 18:2n-6→→→20:4n-6 sequentially use *FADS2*, a usually rapid elongation system, and *FADS1*. Our current results are consistent with hypothesis emerging from our work in lymphoblasts that arachidonic acid would be greater for individuals with the I/I genotype compared to D/D because of basal *FADS1* expression (Reardon, et al. 2012). The immediate precursor 20:3n-6 is lower, and the immediate product, 20:4n-6, is higher, following the hypothesized *FADS1* expression. The precursor 18:2n-6 is not different among genotypes, pointing to the *FADS1* step as the driver for the observed altered levels. Future studies with larger and more diverse samples are warranted to further evaluate the functional implications of this novel indel polymorphism.

Linoleic acid (18:2n-6) is the most prevalent PUFA in the U.S. diet, with mean intake exceeding 13 grams per day (Brenna and Lapillonne 2009; Naqvi, et al. 2012). While it is a precursor of ARA, LA lowering does not reduce circulating ARA because the overwhelming amount of diet LA delivered by seed oils, principally soy oil in the US (Blasbalg, et al. 2011), causes ARA levels to reach a plateau (Gibson, et al. 2013). For instance, reduction of LA intake from 8.5 to 4.2 g/d over 4 weeks resulted in no significant change in plasma PL ARA (9.7 to 9.9%,w/w), though DHA rose 8% (3.47 to 3.77%,w/w) (Wood, et al. 2014). Dietary preformed ARA is required to overcome the maximum and increase circulating ARA (O’Dea and Sinclair 1985). Increasing ARA is accompanied by increasing ARA-derived eicosanoid synthesis including vasoactive species (Ferretti, et al. 1997; Nelson, et al. 1997), though oral ARA consumption at least in the short term do not produce acute clinical effects. How sustained basal ARA levels influence chronic disease progression is not well understood as an isolated factor. The 8% difference in basal ARA probably reflects differences in maximal levels for any particular individual. However, GWAS evidence suggests that the *FADS* gene cluster is associated with diseases that may be of the complex inflammatory type, such as colorectal cancer risk (Zhang, Jia, et al. 2014) and cardiovascular disease (Zhang, Johnson, et al. 2014). The *FADS* gene cluster was found in association with asthma pathogenesis using eQTL mapping (Sharma, et al. 2014), and a *FADS* SNP was found to influence synthesis of arachidonic acid (ARA) and synthesis of pro-inflammatory lipoxygenase products (Hester, et al. 2014).

### Dietary Practices and LCPUFA

Over many generations in India, approximately 35% of the population follows the traditional lacto-vegetarian diet practice (Raheja, et al. 1993; Key, et al. 2006); our Indian cohort followed this trend with responses to dietary pattern questions indicating 38% vegetarian (unpublished), congruent with our dietary instruments applied to research participants in other studies that consistently show a vegetarian dietary pattern and generally low meat consumption in this region (Gadgil, et al. 2014). As plant sources contain only the LCPUFA precursors LA and ALA, individuals on plant based diets must biosynthesize LCPUFA endogenously. The higher frequency of I/I alleles (67.5%) among Indians supports the hypothesis that insertion genotype and its associated haplotype is under selective pressure due to its higher metabolic capacity to convert precursors (LA and ALA) to LCPUFA. This I/I genotype may be favored in populations depending on vegetarian diets and possibly populations having limited access to diets rich in LCPUFA, especially fatty fish.

Homo sapiens evolved eating a diet characterized by 4:1 to 1:1 ratio of n-6/n-3 PUFA (Eaton, et al. 1998). Based on the diet of evolutionary adaptedness (DEA), the overall n-6/n-3 estimate was found to be about 0.79 (Eaton, et al. 1998), whereas, the modern Western diet has undergone an extraordinary increase in n-6 linoleic acid resulting in a ratio above 10/1, and is likely to be even higher in Indian diet, estimate to be 20-50 (Raheja, et al. 1993; Simopoulos 2002). Substantial evidence has accumulated that this dramatic shift is a risk factor for lifestyle-related diseases, such as cardiovascular, diabetes, cancer, and inflammation-related diseases (Raheja, et al. 1993; Simopoulos 2002). Most commercially produced seed oils, sunflower, safflower, peanut, grapeseed, cottonseed and corn, contain very high levels of n-6 LA; individuals with I/I genotype having higher metabolic capacity to convert precursors to longer chain PUFA may be at increased risk for proinflammatory disease states as they efficiently convert LA to ARA. Put another way, individuals with the I/I genotype may be vulnerable to ill-health when adopting a diet rich in n-6 LA which severely reduce synthesis of anti-inflammatory n-3 LCPUFA because n-6 competes with n-3 to access the Δ-6 desaturase enzyme. Moreover, n-6 PUFA compete with and antagonize against incorporation of n-3 PUFA into tissue (Alvheim, et al. 2012), challenging the metabolic requirement for structural n-3 PUFA that are especially concentrated in neural tissue (Diau, et al. 2005).

### Role of Genetic Factors in the Conversion of PUFA Precursors to Products

Metabolism studied using stable isotope labeling, candidate gene SNP, GWAS and metabolomics show interindividual variation in the conversion of PUFA precursors to LCPUFA products depends on genetic factors (Emken, et al. 1994; Schaeffer, et al. 2006; Tanaka, et al. 2009; Illig, et al. 2010). An early FADS gene cluster association study found an inherited component accounting for 28% of the variance in ARA levels among minor allele carriers of a 11 SNP haplotype (Schaeffer, et al. 2006). Minor allele carriers showed lower serum phospholipid ARA. Several independent association studies replicated these findings, showed PUFA levels to be associated with genotypes (Malerba, et al. 2008; Martinelli, et al. 2008; Xie and Innis 2008; Bokor, et al. 2010; Rzehak, et al. 2010; Lattka, et al. 2011; Sergeant, et al. 2012). Our present results show product-precursor differences, arachidonic acid to linoleic acid, to be 13% and 31% greater for I/D and I/I compared to D/D, respectively. Converging data from genetic studies show SNP variants within intron 1 of *FADS2* to be associated with human phenotypes such as IQ scores, blood fatty acid levels and complex diseases (Caspi, et al. 2007; Aulchenko, et al. 2009; Standl, et al. 2011; Steer, et al. 2012; Groen-Blokhuis, et al. 2013; Zhang, Jia, et al. 2014). For instance, commonly reported SNP variants (rs174575, and rs1535) which are associated with increased IQ scores, blood fatty acid levels and complex diseases are <600 bp and <5000 bp upstream from the *FADS2* indel, respectively. In humans, common SNP variants are often found to follow indels (Lu, et al. 2012), suggesting the hypothesis that rs174575 and/or rs1535 are tags for the functional genomic indel that directly modulates binding at the nearby SRE.

### Putative Hypothesis rs66698963 Locus Control Bidirectional Regulation

Our previous cell culture data indicating that *FADS1* and *FADS2* expression *(mRNA)* is upregulated in response to SREBP modulators suggests that this intronic region is a master switch controlling bidirectional regulation of *FADS1* and *FADS2*, as illustrated in supplementary fig. S21. This hypothesis is analogous to regulatory control elsewhere in the human genome: a genome-wide survey of gene organization estimates 11.6% of human genes are bidirectionally oriented (Liu, et al. 2011). For instance, LXR regulated ABC transporters (*ABCG5* and *ABCG8*), which arose by gene duplication events, are arranged in a head-to-head orientation on HSA 2p21. They are transcribed in divergent directions and share common regulatory elements (Remaley, et al. 2002). Similarly, *COL4A1* and *COL4A2* oriented head-to-head on HSA 13q34 also contain a bidirectional transcription unit (Pollner, et al. 1997).

## Conclusions

Collectively our data suggest that *FADS2* indel genotypes contribute to individual variability in response to PUFA consumption. In the vegan or vegetarian scenario in which only, or primarily, precursors would be consumed, tissue LCPUFA would depend on the relative proportions of 18:2n-6 and 18:3n-3. In a food system dominated by linoleic acid, I/I genotype carriers would maintain higher basal arachidonic acid and presumably greater inflammatory potential and attendant higher rates of chronic disease related to inflammation. Balanced consumption of precursors would be particularly important for I/I genotype carriers. I/I carriers consuming excess 18:2n-6 may particularly benefit from consumption of EPA and DHA which bypass the desaturation steps, as a direct balance to ARA. Further, D/D individuals may also benefit from EPA and DHA consumption in pregnancy and in development when neural and other tissue dependent on omega-3 LCPUFA are rapidly develop. For chronic diseases exacerbated by inflammation, D/D carriers maintain lower ARA levels and thus lower inflammatory potential. They are predicted to be less vulnerable to excess 18:2n-6 intake. Future studies should incorporate *FADS2* indel screening as a potential genetic marker for studying LCPUFA regulation.

## Materials and Methods

### Biological Sampling Details

A total of 311 human samples (blood, breastmilk, and placenta) were obtained from Kansas City, Rochester NY, and seven breast milk banks around the U.S. and Canada. Similarly, a total of 234 human blood samples were obtained from Pune City, India. All were used for genotyping. Only RBC (Red Blood Cells) samples from Kansas City were used for fatty acid profiling.

### Study Approvals and Baseline Characteristics

The study was approved by the institutional review boards on human subjects research from all participating institutions, as follows. The University of Kansas Medical Center Human Subjects Committee, University of Rochester Research Subjects Review Board, KEM Hospital Ethics Committee and Institutional Review Board of Sinhgad Institute, India, all approved human sample collection and written informed consent was obtained from all donors. The Cornell University Institutional Review Board approved collection of samples and waived consent because the samples were collected and anonymized at human milk banks (exemption 4), as follows: Mothers’ Milk Bank, San Jose, CA; Bronson Mothers’ Milk Bank, Kalamazoo, MI; Mothers’ Milk Bank, a program of Rocky Mountain Children’s Health Foundation, Denver, CO; Ohio Health Mothers’ Milk Bank, Columbus, OH; Mothers’ Milk Bank at Austin, Austin, TX; Mothers’ Milk Bank of Northeast, Newton Upper Falls, MA; British Columbia Women’s Milk Bank, Vancouver, BC, Canada. Subjects (n=234) from Indian cohort were all Asian Indians, among them n=122 were women (ages 18 to 43 years) and n=112 were men (ages 23 to 36 years). Dietary patterns from surveys (not presented) indicated that n=89 (38.03%) were vegetarians and n=145 (61.97%) were omnivores. The US cohort (311 participants) were all females. Breastmilk (n=69) samples from US cohort are banked samples. Placental (n=41) samples are from pregnant adolescents (ages ≤18 years) who were recruited from the Rochester Adolescent Maternity Program in Rochester, NY (Cao, et al. 2014). Detailed demographic characteristics of Kansas cohort (n=201) are provided elsewhere (Carlson, et al. 2013). All participants were English speaking and are between ages 16 to 36 years. RBC fatty acid analysis were available only for the Kansas City participants (n=199). Fatty acid data is not available from the Indian cohort.

### Genotyping rs66698963

Genomic DNA was extracted and was used to amplify a 629 base pair fragment flanking the SRE within intron 1 of FADS2 (GenBank Accession# NT_167190.1) by PCR as previously described (Reardon, et al. 2012). The following primer pairs were used to amplify 629 bp fragment: FADS2 forward primer 5′ TTTCTCAAAGGCCGTGGTGT 3′, FADS2 reverse primer 5′ AGTGCTAACCACTCCTGGAA 3′. The amplified products were run on 2% agarose gels until the two alleles, if present, were well separated. The gels were stained with ethidium bromide and visualized under UV to identify indel genotypes. A gel demonstrating detection of the three possible genotypes for the 22 bp indel is presented in Figure 1. Samples were assigned pairs of genotypes of I (insertion) or D (deletion) as I/I, I/D, or D/D.

### Genotype Frequency of rs66698963 in 1000GP

VCF files of genetic variations data from the 1000GP (phase 3) were downloaded from the official FTT site (ftp://ftp.1000genomes.ebi.ac.uk/vol1/ftp/release/20130502/). The indel rs66698963 was not directly annotated in the VCF file while another indel rs373263659 with the same length of insertion was present at the same region. Using Sanger sequencing results from our previous study (Reardon, et al. 2012), we determined that rs373263659 is a misannotation of rs66698963. To confirm this observation and to further test the accuracy of indel calling in 1000GP, we examined the read mapping results for 16 Japanese individuals, for whom we have Sanger sequencing results. BAM files for these 16 individuals were downloaded from each individual’s directory on the FTP site and visualized with SAMtools (Li, et al. 2009). After confirming that rs373263659 is a misannotation of rs66698963 and the 1000GP has acceptable calling accuracy for this indel, we used directly the frequency of rs373263659 as that of rs66698963.

In the 1000GP, there are in total 2504 individuals from five continental regions and 26 global populations. Four admixed American populations were excluded from our analysis because of their admixture history. After the exclusion, there are seven populations of African ancestry (AFR): Yoruba in Ibadan, Nigeria (YRI), Luhya in Webuye, Kenya (LWK), Gambian in Western Divisions in the Gambia (GWD), Mende in Sierra Leone (MSL), Esan in Nigeria (ESN), Americans of African Ancestry in SW USA (ASW), African Caribbeans in Barbados (ACB); five populations of European ancestry (EUR): Utah Residents with Northern and Western European Ancestry (CEU), Toscani in Italia (TSI), Finnish in Finland (FIN), British in England and Scotland (GBR), Iberian Population in Spain (IBS); five populations of East Asian ancestry (EAS): Han Chinese in Beijing, China (CHB), Japanese in Tokyo, Japan (JPT), Southern Han Chinese (CHS), Chinese Dai in Xishuangbanna, China (CDX), Kinh in Ho Chi Minh City, Vietnam (KHV); and five populations of South Asian ancestry (SAS): Gujarati Indian from Houston, Texas (GIH), Punjabi from Lahore, Pakistan (PJL), Bengali from Bangladesh (BEB), Sri Lankan Tamil from the UK (STU), Indian Telugu from the UK (ITU).

### Testing for Positive Selection on the FADS Region

SFS-based statistics, including genetic diversity (_TT_) Tajima’s D (Tajima 1989), and Fay and Wu’s H (Fay and Wu 2000), and population differentiation-based statistics, F_*ST*_ (Weir and Cockerham 1984) and population branch statistics (PBS) (Yi, et al. 2010) were calculated using in-house Perl scripts, which are available upon request. F_*ST*_ and PBS were calculated for each genetic variant. Genetic diversity, Tajima’s D, and Fay and Wu’s H were calculated using a sliding-window approach with window size of 5 Kb and moving step of 1 Kb. Statistical significance for these statistics were assessed using the ranking of genome-wide variants or windows, respectively. For example, if the Tajima’s D value for a specific 5-Kb window is among the top 5% of all genome-wide windows, then it is considered as statistically significant with an empirical *p* value <0.05. PBS was estimated based on pair-wise F_*ST*_ values among three continental regions (Yi, et al. 2010). For a specific locus, the PBS value for a population represents the amount of allele frequency change in the history of this population since its divergence from the other two populations. The PBS estimate for a population depends on the choice of the other two populations. Therefore, to calculate PBS for South Asians, we chose three combinations of continental regions: AFR, EUR and SAS; AFR, EAS and SAS; EUR, EAS and SAS.

Haplotype-based tests for positive selection, including the Integrated Haplotype Score (iHS) (Voight, et al. 2006), the number of segregating sites by length (nSL) (Ferrer-Admetlla, et al. 2014), and Cross-population Extended Haplotype Homozygosity (XPEHH) (Sabeti, et al. 2007), were calculated using the software selscan v 1.1.0a (Szpiech and Hernandez 2014). All SNPs with minor allele frequency greater than 5% in at least one of the four continental regions were included in our calculation. Genetic variants without information on their ancestral allele were excluded from analyses. Information on ancestral allele was retrieved directly from the VCF file with an exception for the indel under study, rs66698963, whose ancestral allele was not defined in the VCF file. Alignment of homologous sequences from human, Chimpanzee and Orangutan showed that neither the insertion nor the deletion allele of rs66698963 is present in Chimpanzee or Orangutan (supplementary fig. S22A). By examining the mapping results of short sequencing reads from a Neanderthal from the Altai Mountains (Prufer, et al. 2014) and an archaic Denisovan individual (Meyer, et al. 2012) to the human genome, we found that the insertion allele is present in both Neanderthal and Denisovan genomes and it is therefore considered as the ancestral allele. Empirical *p* values were inferred based on the genome-wide distribution of the statistics. The haplotype bifurcation diagrams and EHH plots were drawn using an R package, *rehh* (Gautier and Vitalis 2012).

### Fatty Acid Analysis

RBC phospholipids (PL) fatty acids were analyzed for 199 samples from Kansas according to methods reported elsewhere (Carlson, et al. 2013) and all fatty acids were correlated with genotype. Briefly, lipids from RBC were extracted according to the modified Folch method (Folch, et al. 1957). Extracted RBC lipids were fractionated by loading on to the TLC plate (Zail and Pickering 1979). Fatty acid methyl esters (FAME) from RBC PL were prepared using boron trifluoride-methanol (Morrison and Smith 1964), and were separated by using an SP-2560 capillary column (100 m; Sigma Aldrich) (Smuts, et al. 2003). Peak integration and analysis was carried out by using a Star 6.41 Chromatography Workstation. Individual peaks were identified by comparing their retention times with those of known standards (PUFA 1 and PUFA 2; Sigma Aldrich), and an equal weight FAME mixture (Supelco 37 Component FAME mix; Sigma Aldrich) was used to adjust fatty acids for area/weight to calculate a final percentage weight of total fatty acids (Carlson, et al. 2013). Fatty acids are expressed as percent weight-for-weight (%,w/w). Percent changes are expressed as Ax/x/AD/D, where A denotes a fatty acid, x = I or D, and D/D is the deletion/deletion genotype. Pairwise statistical analysis was conducted using Student’s t-test at P<0.05 and data are presented as mean ± 95% confidence interval. Secondarily, the non-parametric Wilcoxon method was also performed to verify significance in case of skewed fatty acid distributions.

## Acknowledgements

This work was supported by NIH grant R01 AT007003 from the National Center for Complementary and Integrative Health (NCCIH) (formerly the National Center for Complementary and Alternative Medicine (NCCAM)) and the Office of Dietary Supplements (ODS), NIH grant R01 HD047315, USDA-NIFA National Research Initiative 2005-35200-15218, and NIH grant R01 GM108805. Its contents are solely the responsibility of the authors and do not necessarily represent the official views of the NIH, USDA, or any of their units. The authors thank Dr. R.R. Ran-Ressler, Cornell University, for assistance with breast milk samples.

---

**Supplementary Table S1.** Genotype Frequencies of rs66698963 in 22 global populations

| Continental Regions | Populations | I/I | I/D | D/D | I | D |
| --- | --- | --- | --- | --- | --- | --- |
| AFR | YRI | 0.53 (57) | 0.38 (41) | 0.09 (10) | 0.72 | 0.28 |
|  | LWK | 0.59 (58) | 0.36 (36) | 0.05 (5) | 0.77 | 0.23 |
|  | GWD | 0.60 (68) | 0.38 (43) | 0.02 (2) | 0.79 | 0.21 |
|  | MSL | 0.55 (47) | 0.38 (32) | 0.07 (6) | 0.74 | 0.26 |
|  | ESN | 0.54 (53) | 0.42 (42) | 0.04 (4) | 0.75 | 0.25 |
|  | ASW | 0.39 (24) | 0.57 (35) | 0.03 (2) | 0.68 | 0.32 |
|  | ACB | 0.48 (46) | 0.47 (45) | 0.05 (5) | 0.71 | 0.29 |
| EWR | CEU | 0.21 (21) | 0.64 (63) | 0.15 (15) | 0.53 | 0.47 |
|  | TSI | 0.20 (21) | 0.60 (64) | 0.21 (22) | 0.50 | 0.50 |
|  | FIN | 0.15 (15) | 0.52 (51) | 0.33 (33) | 0.41 | 0.59 |
|  | GBR | 0.12 (11) | 0.58 (53) | 0.30 (27) | 0.41 | 0.59 |
|  | IBS | 0.14 (15) | 0.59 (63) | 0.27 (29) | 0.43 | 0.57 |
| EAS | CHB | 0.48 (49) | 0.45 (46) | 0.08 (8) | 0.70 | 0.30 |
|  | JPT | 0.54 (56) | 0.43 (45) | 0.03 (3) | 0.75 | 0.25 |
|  | CHS | 0.22 (23) | 0.62 (65) | 0.16 (17) | 0.53 | 0.47 |
|  | CDX | 0.10 (9) | 0.58 (54) | 0.32 (30) | 0.39 | 0.61 |
|  | KHV | 0.08 (8) | 0.68 (67) | 0.24 (24) | 0.42 | 0.58 |
| SAS | GIH | 0.80 (82) | 0.19 (20) | 0.01 (1) | 0.89 | 0.11 |
|  | PJL | 0.65 (62) | 0.29 (28) | 0.06 (6) | 0.79 | 0.21 |
|  | BEB | 0.59 (51) | 0.40 (34) | 0.01 (1) | 0.79 | 0.21 |
|  | STU | 0.66 (67) | 0.33 (34) | 0.01 (1) | 0.82 | 0.18 |
|  | ITU | 0.78 (80) | 0.22 (22) | 0.00 (0) | 0.89 | 0.11 |

NOTE: 1. The numbers of individuals of specific genotype are presented in parenthesis.

2. Consult methods section for populations represented by the population codes.

**Supplementary Table S2.** Kansas cohort participant’s characteristics (n=199)

| Genotype | I/I | I/D | D/D |
| --- | --- | --- | --- |
| Caucasian Age | 27.43 ± 4.01 (n=13) | 26.45 ± 4.66 (n=49) | 26.82 ± 4.33 (n=76) |
| Caucasian BMI | 24.94 ± 4.68 (n=13) | 26.76 ± 4.99 (n=49) | 26.12 ± 4.87 (n=76) |
| African American Age | 24.2 ± 5.79 (n=20) | 25.19 ± 5.37 (n=27) | 23.40 ± 3.24 (n=12) |
| African American BMI | 28.61 ± 4.41 (n=20) | 30.2 ± 4.08 (n=27) | 28.91 ± 7.05 (n=12) |
| Others Age | 35 (n=1) | None | 29 (n=1) |
| Others BMI | 26.3 (n=1) | None | 34.55 (n=1) |

**Supplementary Figure S1.**
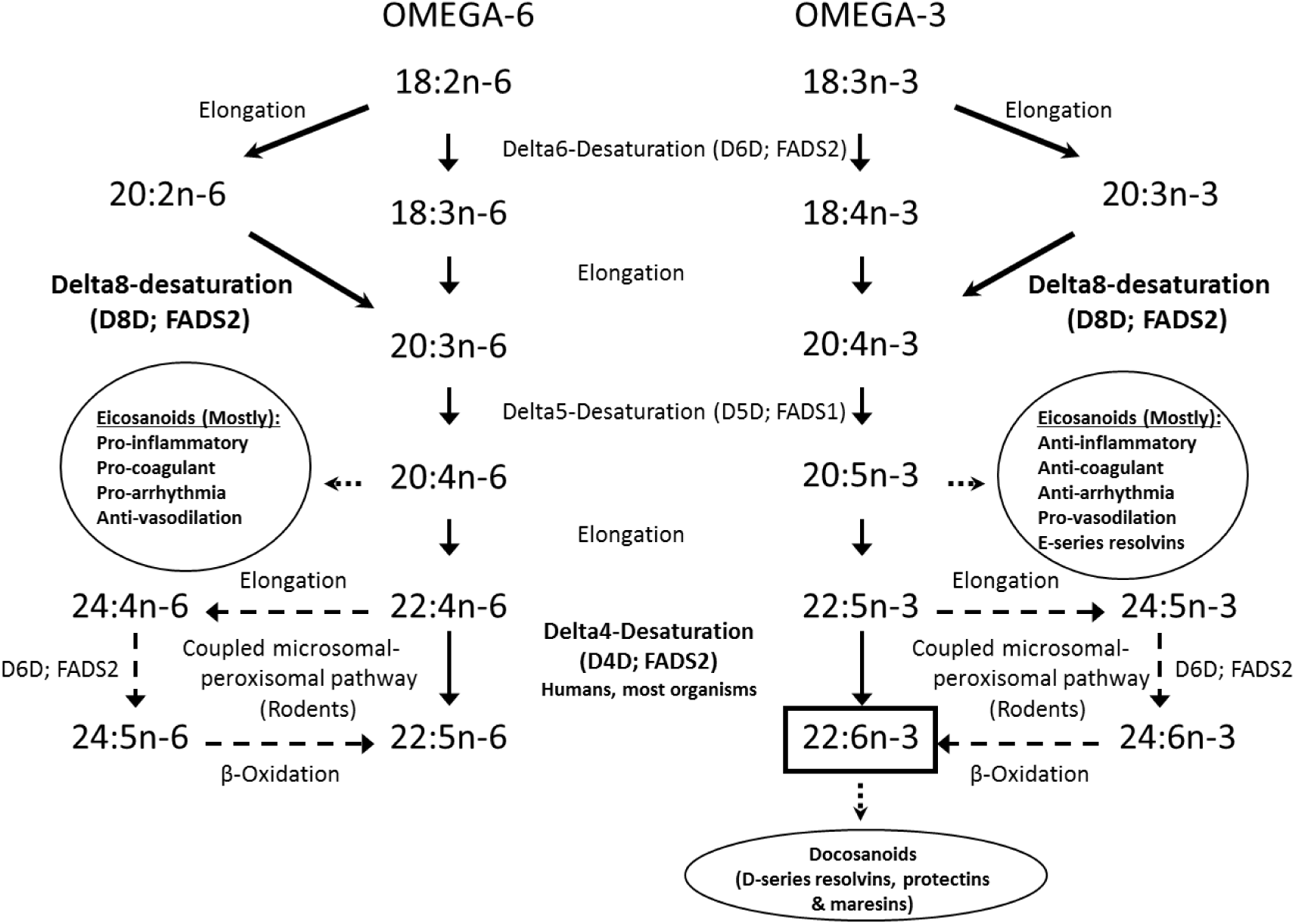
Both n-3 and n-6 LCPUFA pathways compete for the same FADS2, FADS1 and Elongation enzymes. The Δ6-desaturase (FADS2) catalyzes critical rate limiting step operating on both 18:3n-3 and 18:2n-6, resulting in the synthesis of 6,9,12,15-18:4 and 6,9,12-18:3 (gamma-linolenic acid), respectively. Δ6-desaturation is followed by elongation to 8,11,14,17-20:4 and 8,11,14-20:3 (dihomo-gamma-linolenic acid) and a rapid Δ5-desaturation (FADS1) to produce biologically active eicosanoid precursors 20:5n-3 (EPA) and 20:4n-6 (ARA), respectively. EPA and ARA can be further elongated and desaturated to yield 22:6n-3 (DHA) and 22:5n-6 (DPA), respectively by the pathway shown, which is accepted as coupled microsomal-peroxisomal pathway, or via a Δ4-desaturase (FADS2).

**Supplementary Figure S2.**
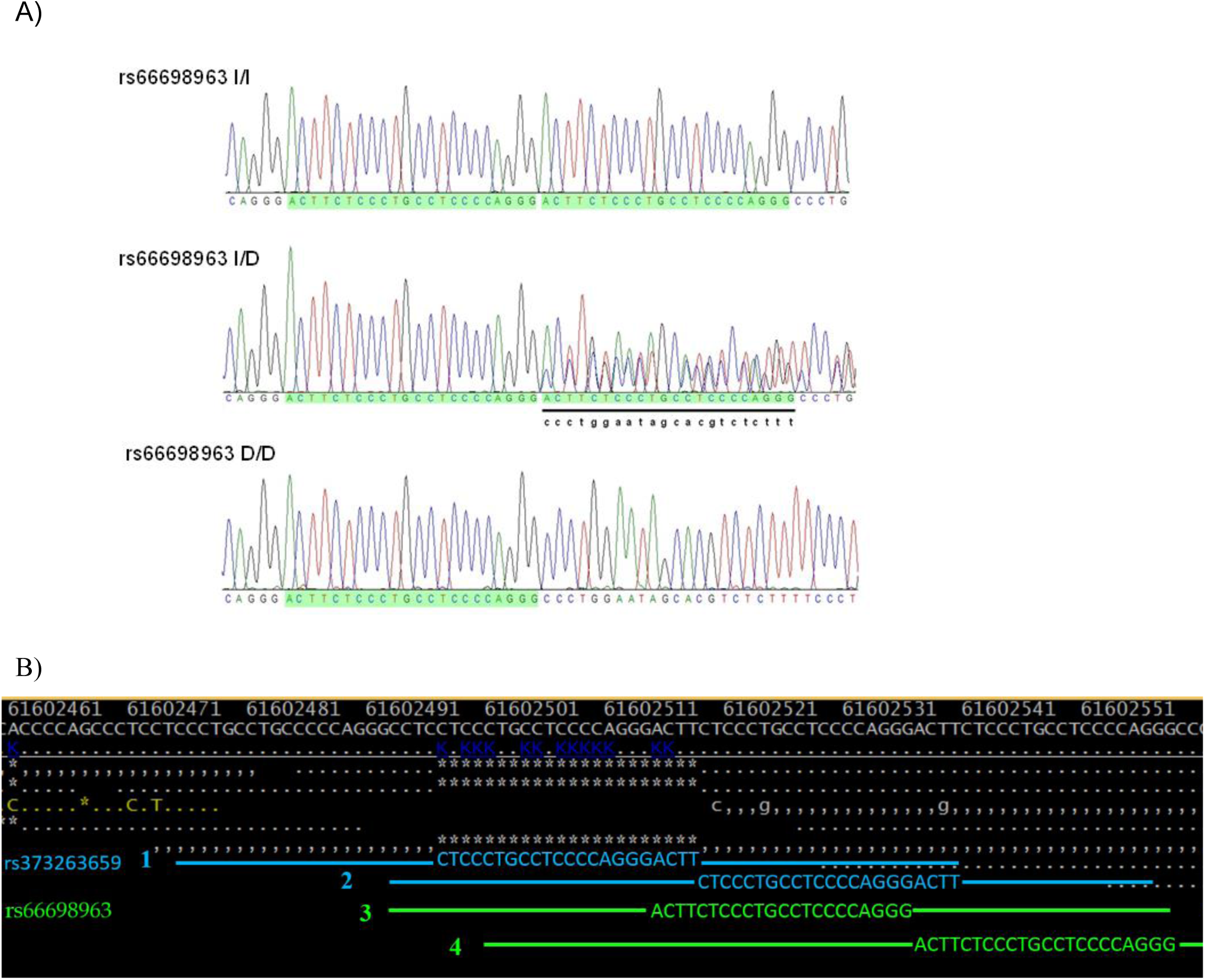
A) Sanger sequencing chromatograms of the 22 bp insertion-deletion (Indel) rs66698963 polymorphism in the Intron 1 of *FADS2;* B) Sequencing reads and mapping results surrounding the rs66698963 using individual HG03838 as an example. In the 1000GP data, there is no annotation for rs66698963. Instead, there is another Indel with dbSNP ID rs373263659. The Insertion allele for rs373263659 is also 22bp long but is slightly different from that for rs66698963. Two tandem 22 bp repeats, 1 and 2 (blue color) represents rs373263659 and 3 and 4 (green color) represents rs66698963. Two possible Indels, rs373263659 and rs66698963, are difficult to differentiate with limited sequencing coverage and short sequencing read lengths. However, they could be differentiated with ease by Sanger sequencing, which has much longer read lengths. Based on our Sanger sequencing results, we confirmed that the correct annotation for this Indel should be rs66698963.

**Supplementary Figure S3.**
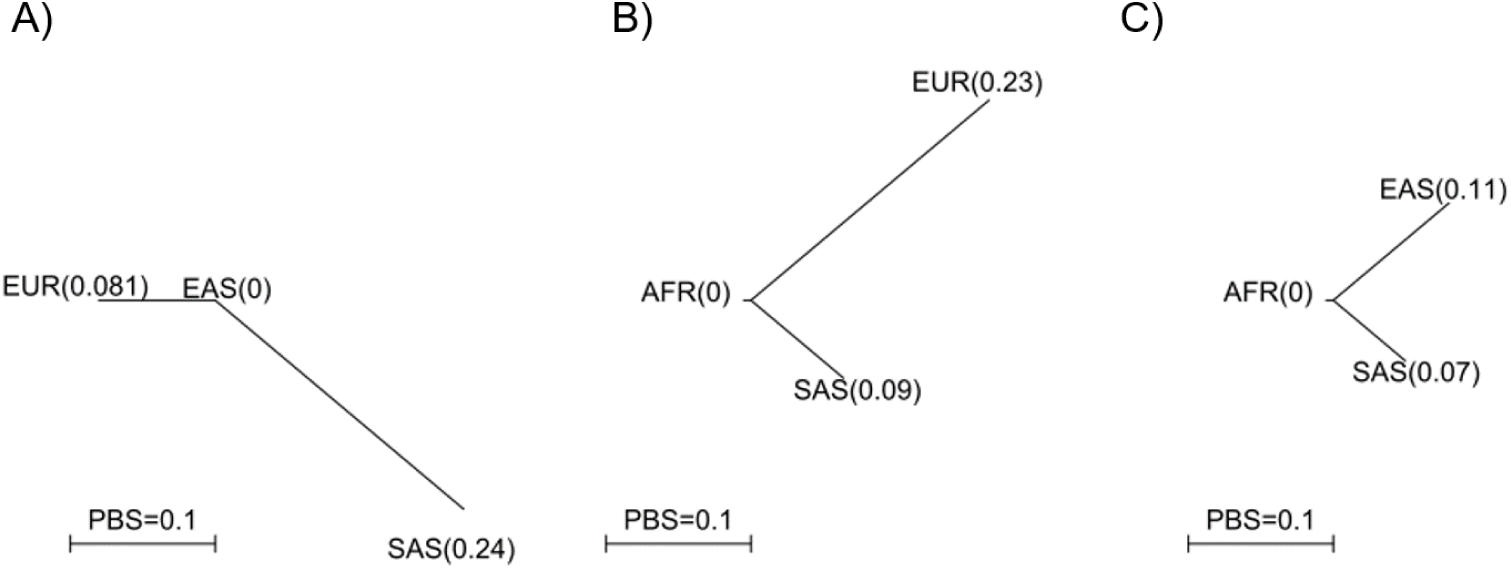
Patterns of Population Branch Statistics (PBS) calculated under three combinations of continental populations.

**Supplementary Figure S4.**
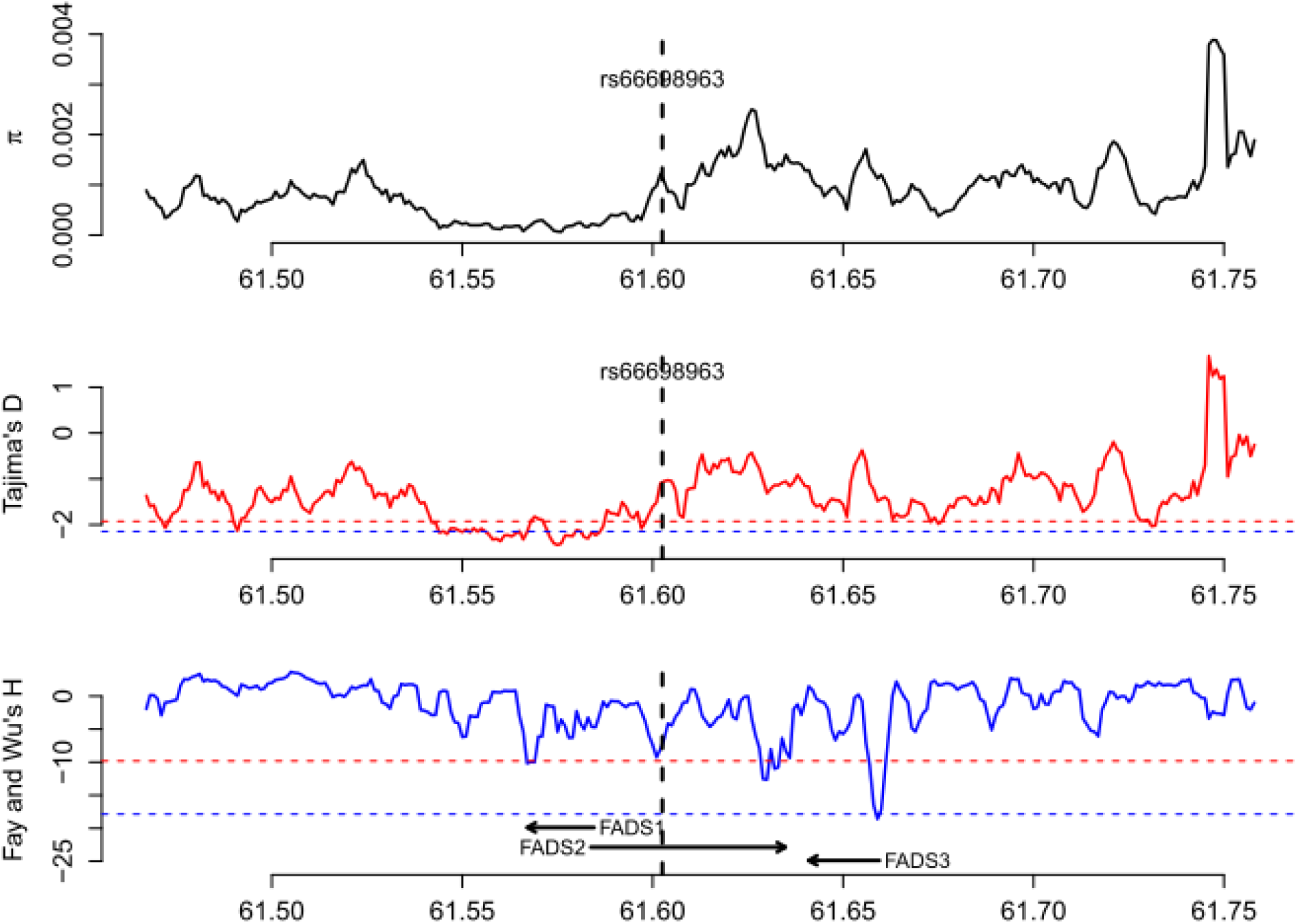
Patterns of genetic diversity (π), Tajima’s D, and Fay and Wu’s H in all populations of African origin (AFR). The red dashed line indicates the 5% genome-wide significance cutoff while the blue dashed line indicates the 1% genome-wide significance cutoff.

**Supplementary Figure S5.**
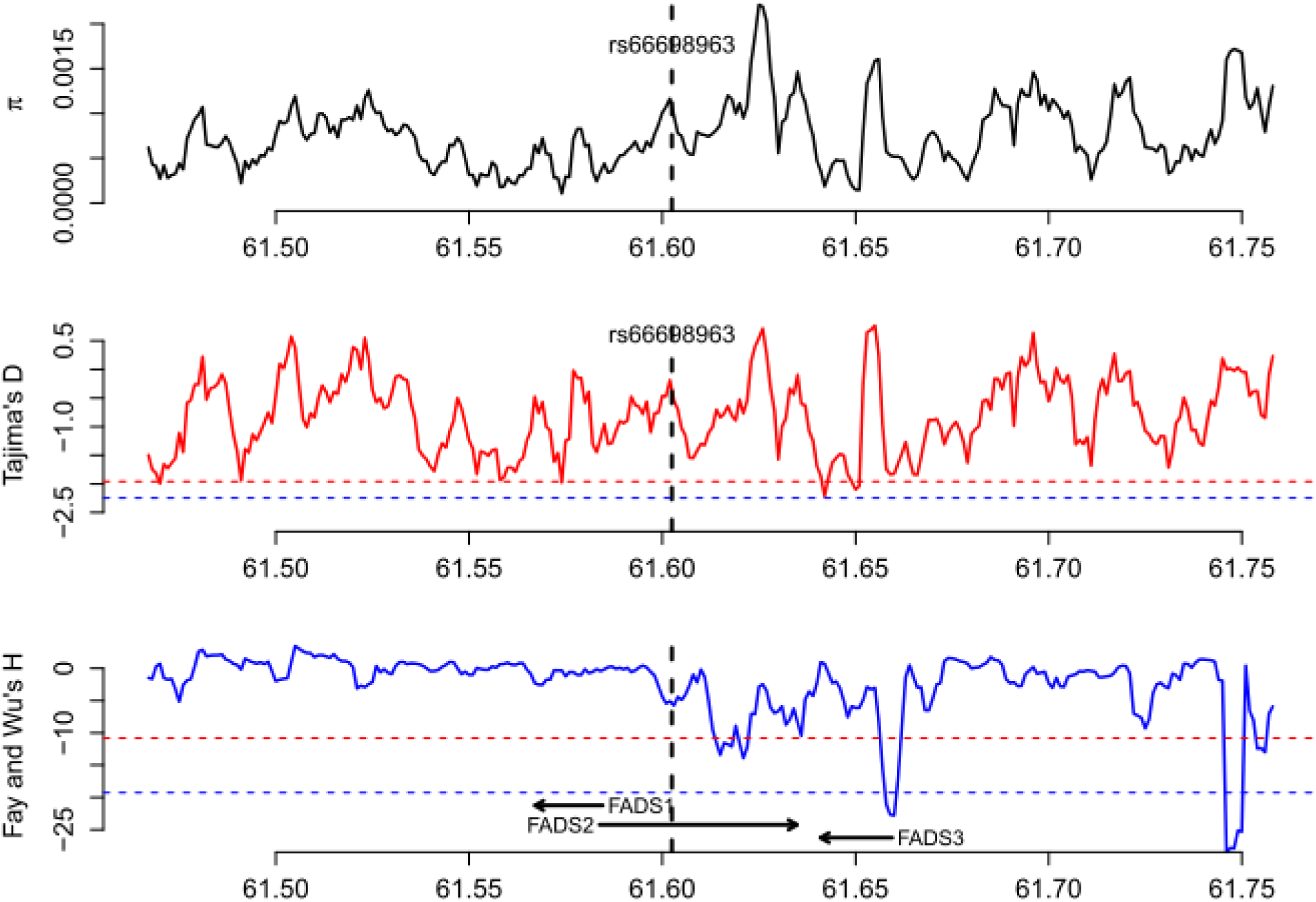
Patterns of genetic diversity (π), Tajima’s D, and Fay and Wu’s H in all populations of European origin (EUR). The red dashed line indicates the 5% genome-wide significance cutoff while the blue dashed line indicates the 1% genome-wide significance cutoff.

**Supplementary Figure S6.**
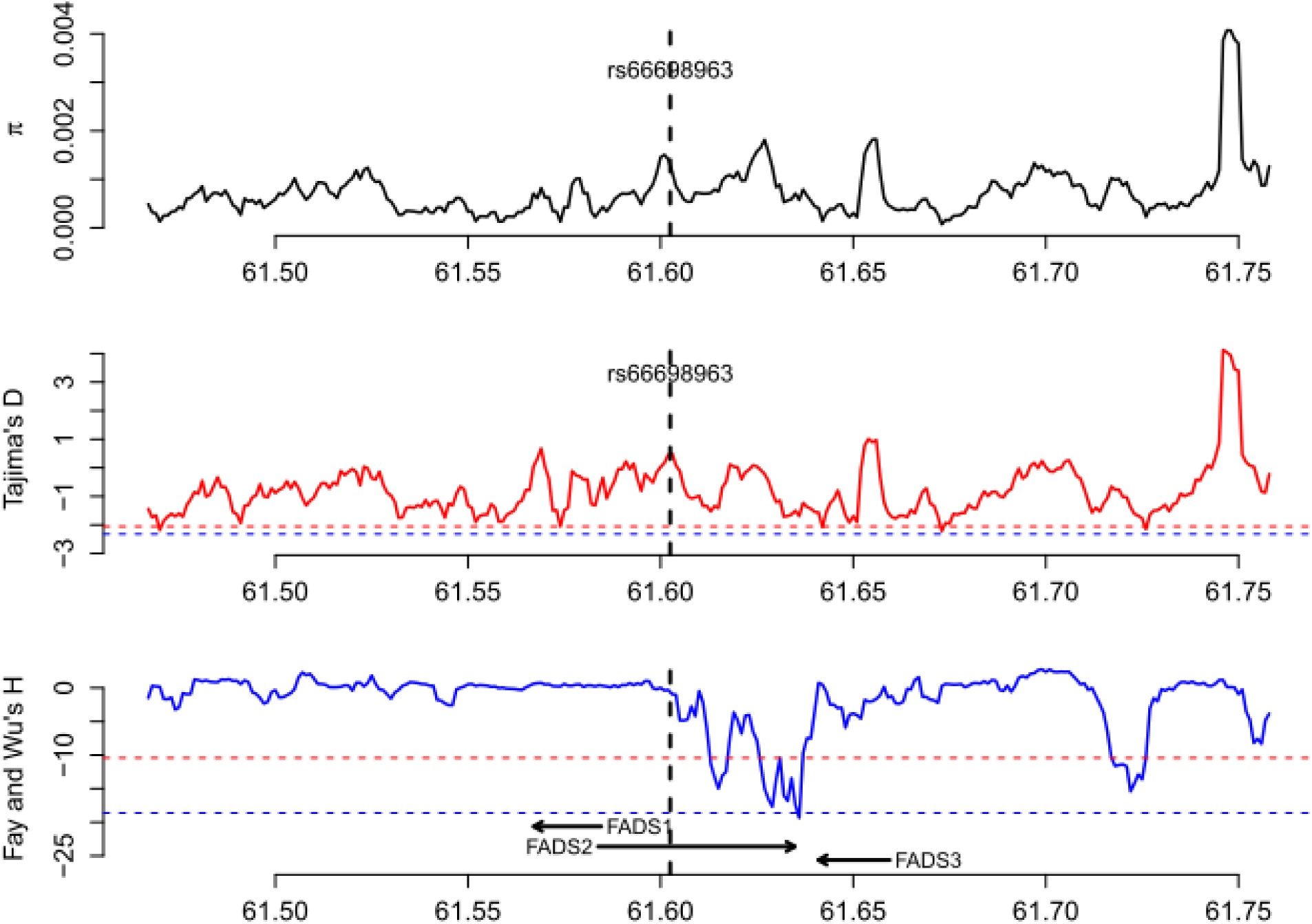
Patterns of genetic diversity (π), Tajima’s D, and Fay and Wu’s H in all populations of East Asian origin (EAS). The red dashed line indicates the 5% genome-wide significance cutoff while the blue dashed line indicates the 1% genome-wide significance cutoff.

**Supplementary Figure S7.**
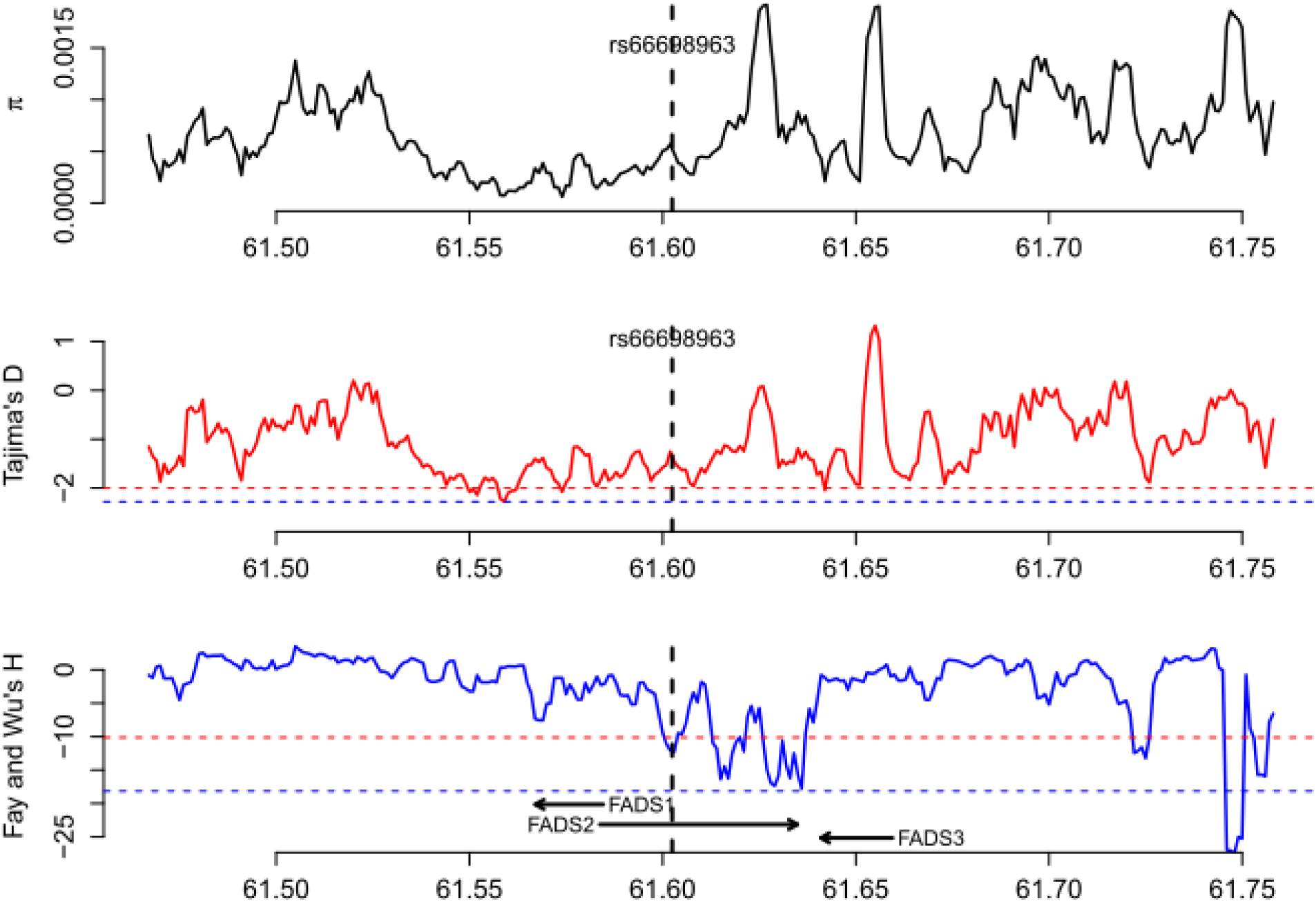
Patterns of genetic diversity (π), Tajima’s D, and Fay and Wu’s H in all populations of South Asian origin (SAS). The red dashed line indicates the 5% genome-wide significance cutoff while the blue dashed line indicates the 1% genome-wide significance cutoff.

**Supplementary Figure S8.**
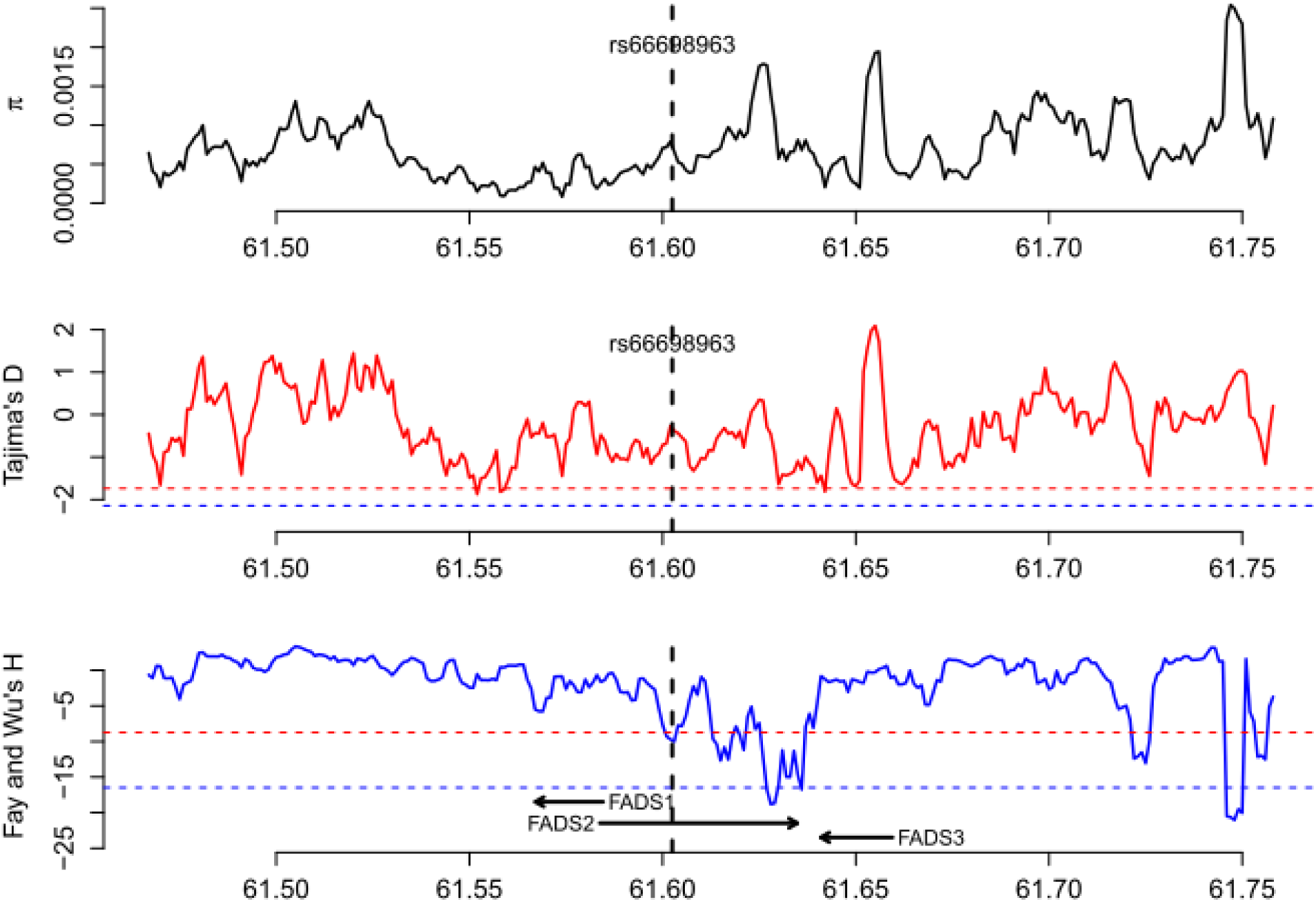
Patterns of genetic diversity (π), Tajima’s D, and Fay and Wu’s H in Bengali from Bangladesh (BEB). The red dashed line indicates the 5% genome-wide significance cutoff while the blue dashed line indicates the 1% genome-wide significance cutoff.

**Supplementary Figure S9.**
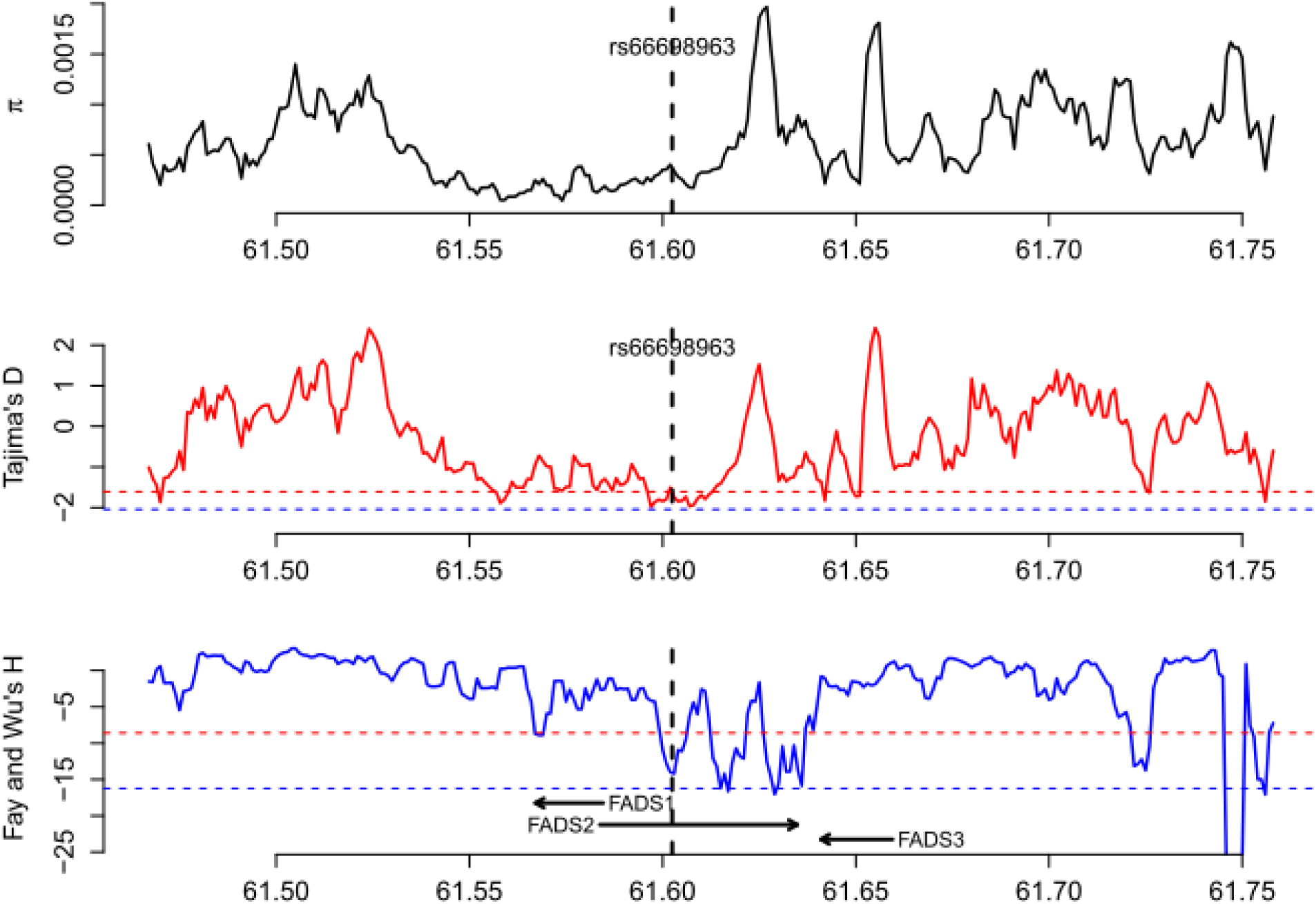
Patterns of genetic diversity (π), Tajima’s D, and Fay and Wu’s H in Gujarati Indian from Houston, Texas (GIH). The red dashed line indicates the 5% genome-wide significance cutoff while the blue dashed line indicates the 1% genome-wide significance cutoff.

**Supplementary Figure S10.**
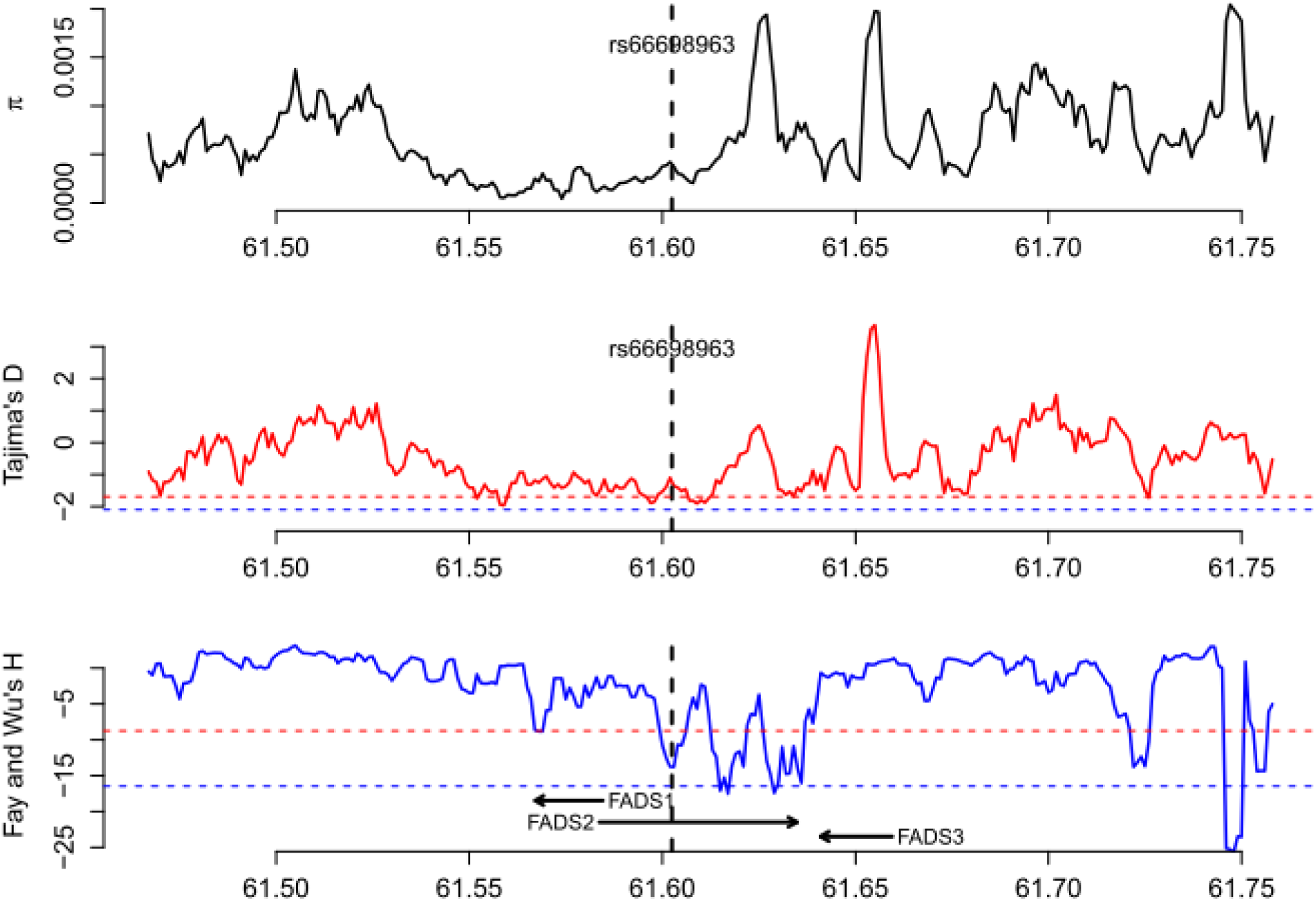
Patterns of genetic diversity (π), Tajima’s D, and Fay and Wu’s H in Indian Telugu from the UK (ITU). The red dashed line indicates the 5% genome-wide significance cutoff while the blue dashed line indicates the 1% genome-wide significance cutoff.

**Supplementary Figure S11.**
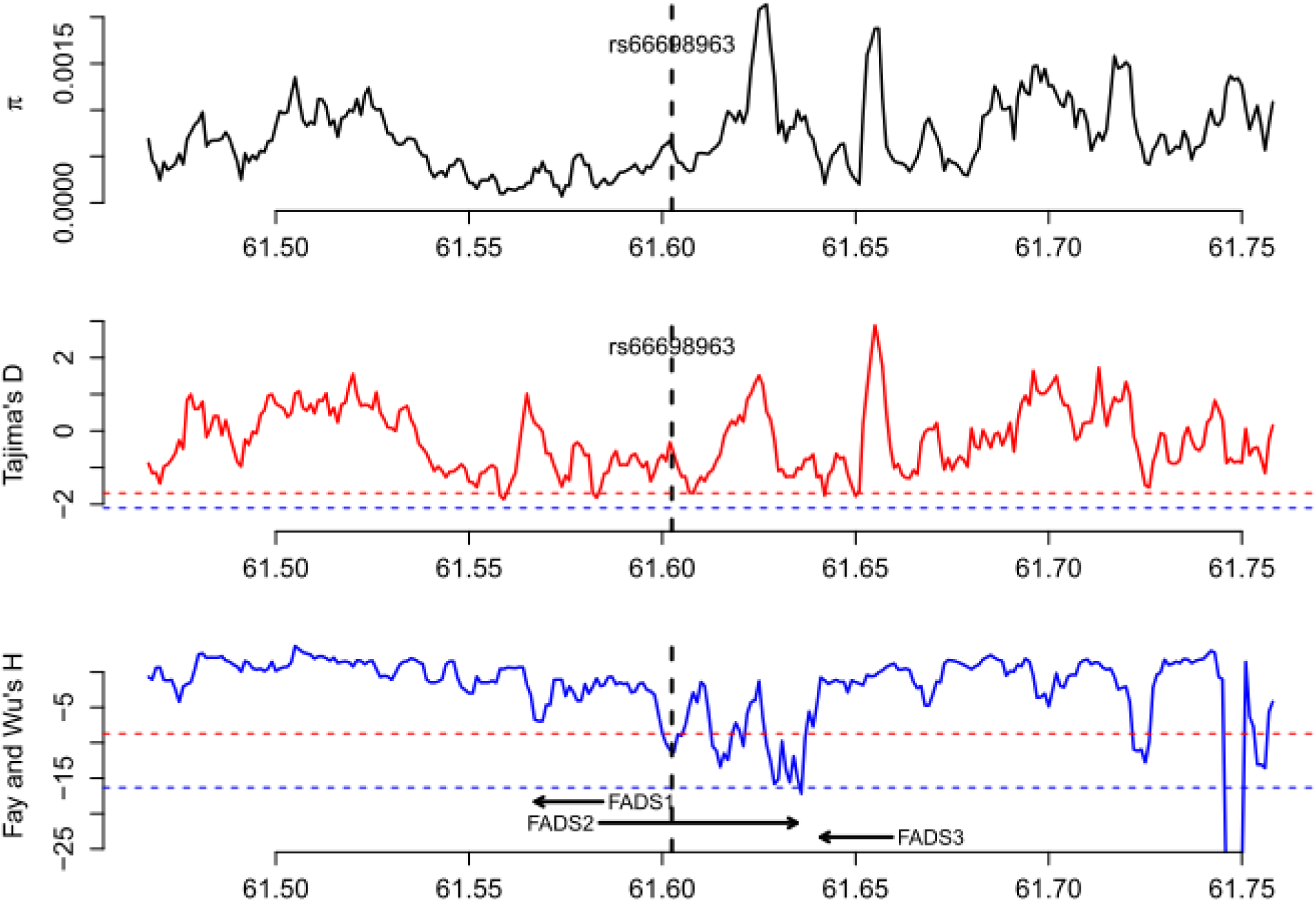
Patterns of genetic diversity (π), Tajima’s D, and Fay and Wu’s H in Punjabi from Lahore, Pakistan (PJL). The red dashed line indicates the 5% genome-wide significance cutoff while the blue dashed line indicates the 1% genome-wide significance cutoff.

**Supplementary Figure S12.**
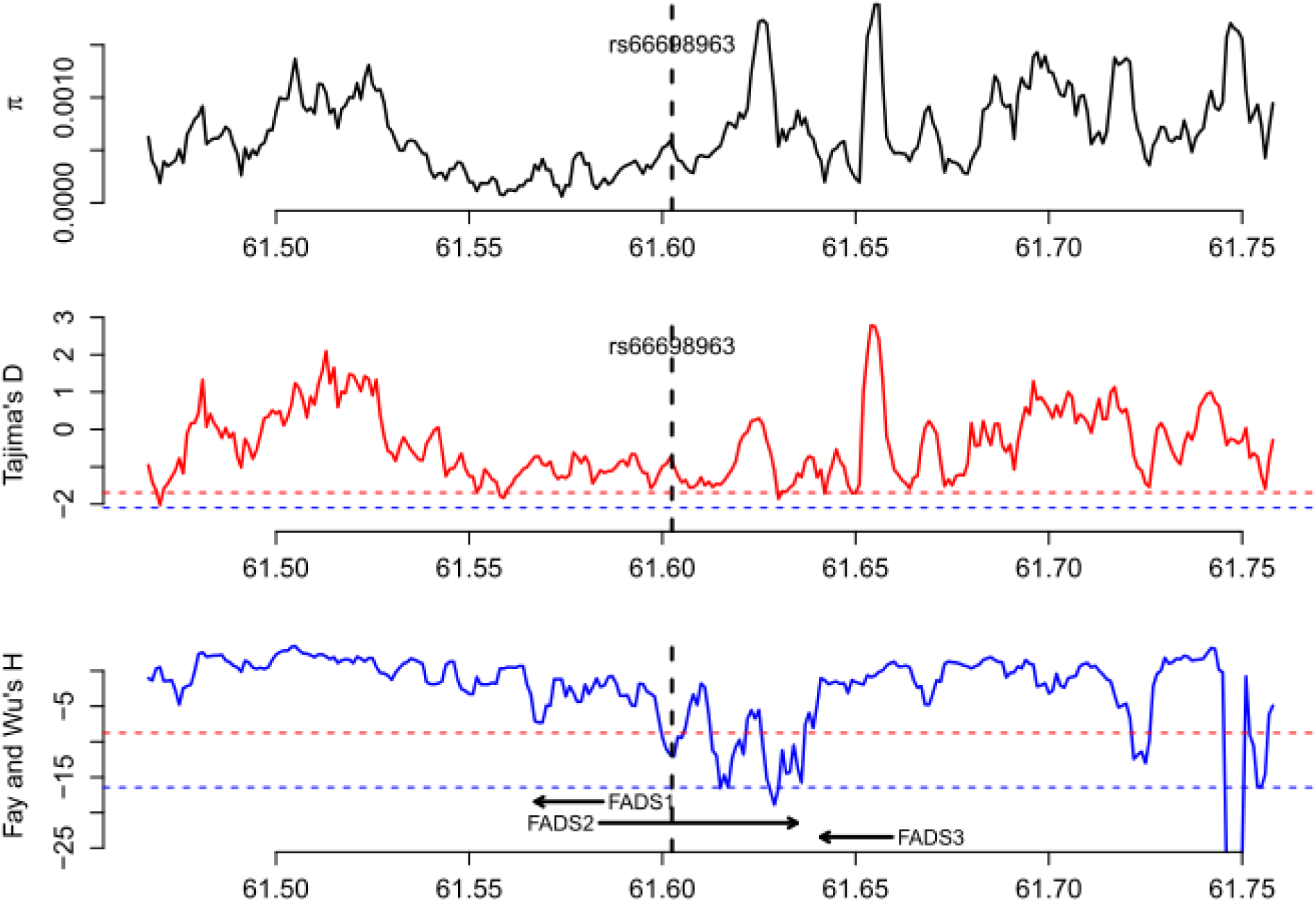
Patterns of genetic diversity (π), Tajima’s D, and Fay and Wu’s H in Sri Lankan Tamil from the UK (STU). The red dashed line indicates the 5% genome-wide significance cutoff while the blue dashed line indicates the 1% genome-wide significance cutoff.

**Supplementary Figure S13.**
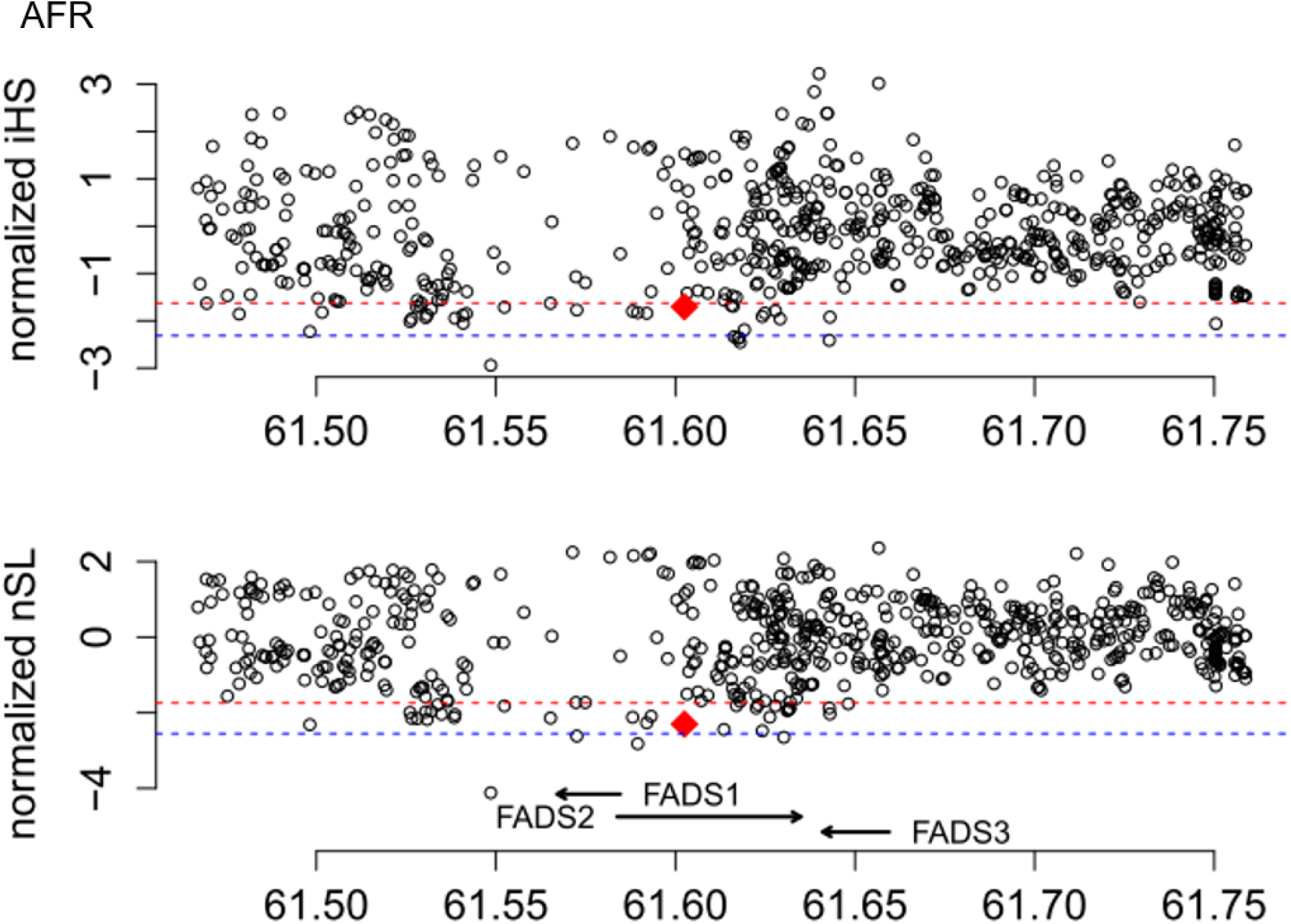
Patterns of normalized iHS and nSL across a 300 Kb region around FADS genes in all seven populations of African origin (AFR). The red dashed line indicates the 5% genome-wide significance cutoff while the blue dashed line indicates the 1% genome-wide significance cutoff.

**Supplementary Figure S14.**
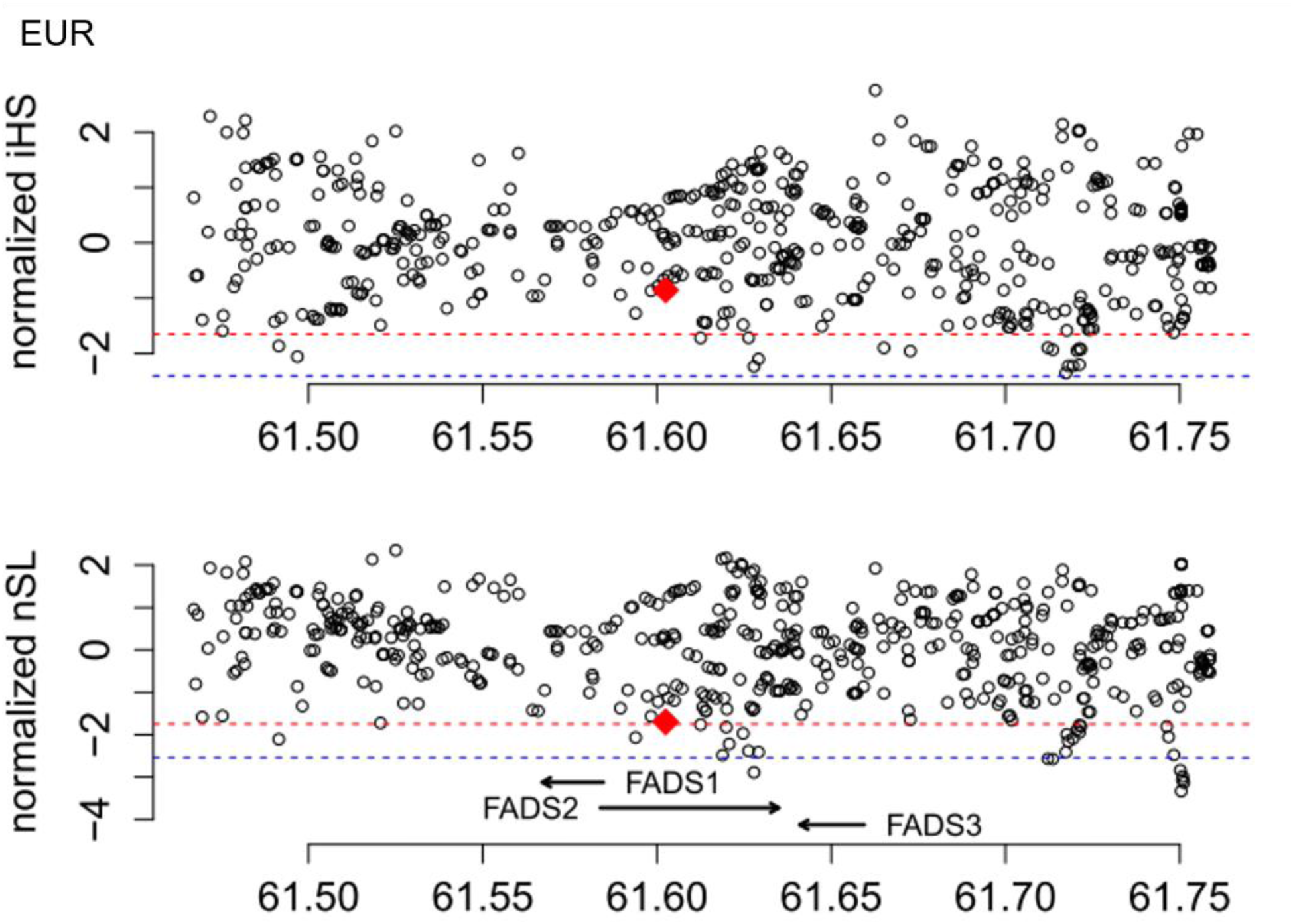
Patterns of normalized iHS and nSL across a 300 Kb region around FADS genes in all five populations of European origin (EUR). The red dashed line indicates the 5% genome-wide significance cutoff while the blue dashed line indicates the 1% genome-wide significance cutoff.

**Supplementary Figure S15.**
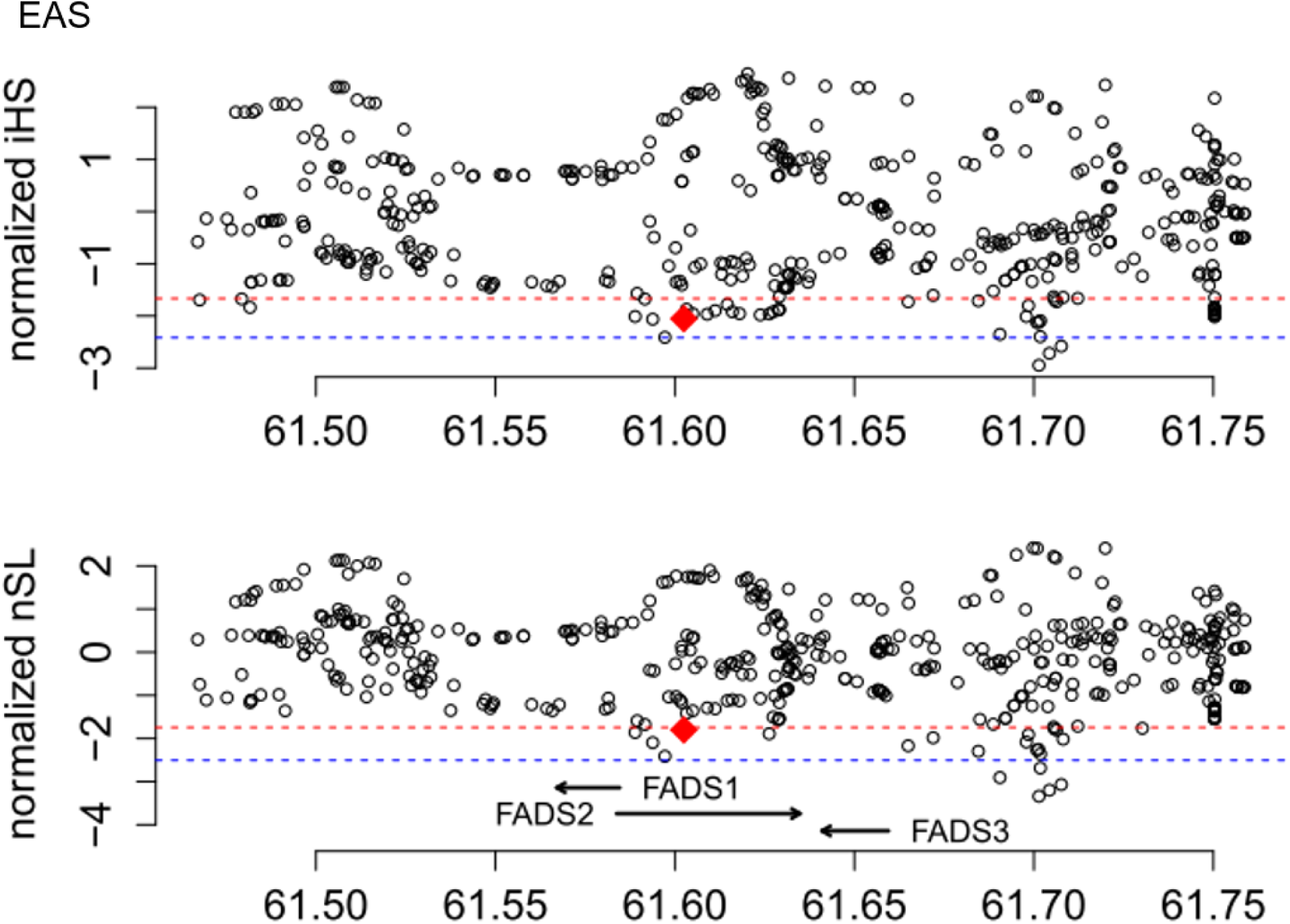
Patterns of normalized iHS and nSL across a 300 Kb region around FADS genes in all five populations of East Asian origin (EAS). The red dashed line indicates the 5% genome-wide significance cutoff while the blue dashed line indicates the 1% genome-wide significance cutoff.

**Supplementary Figure S16.**
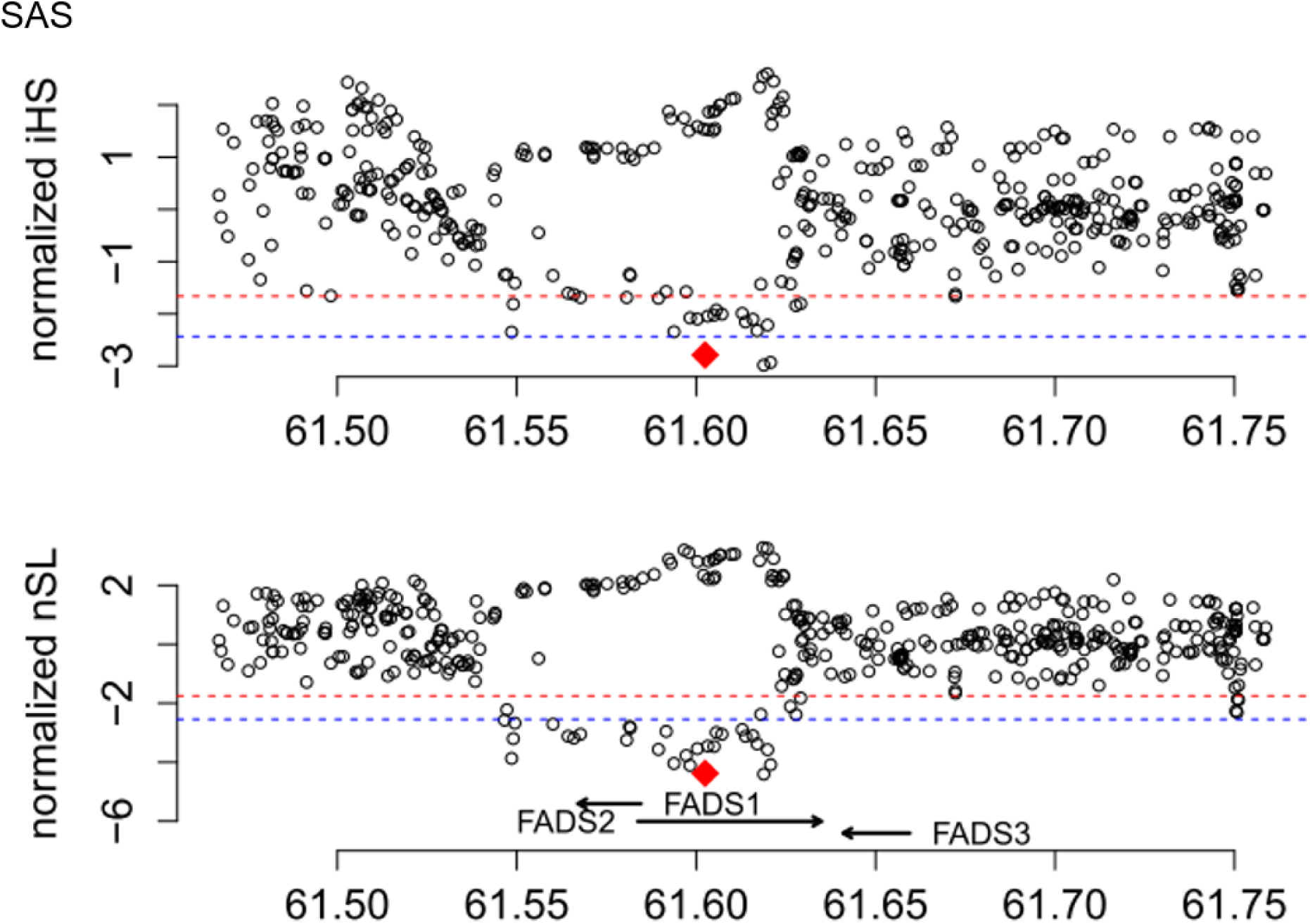
Patterns of normalized iHS and nSL across a 300 Kb region around FADS genes in all five populations of South Asian origin (SAS). The red dashed line indicates the 5% genome-wide significance cutoff while the blue dashed line indicates the 1% genome-wide significance cutoff.

**Supplementary Figure S17.**
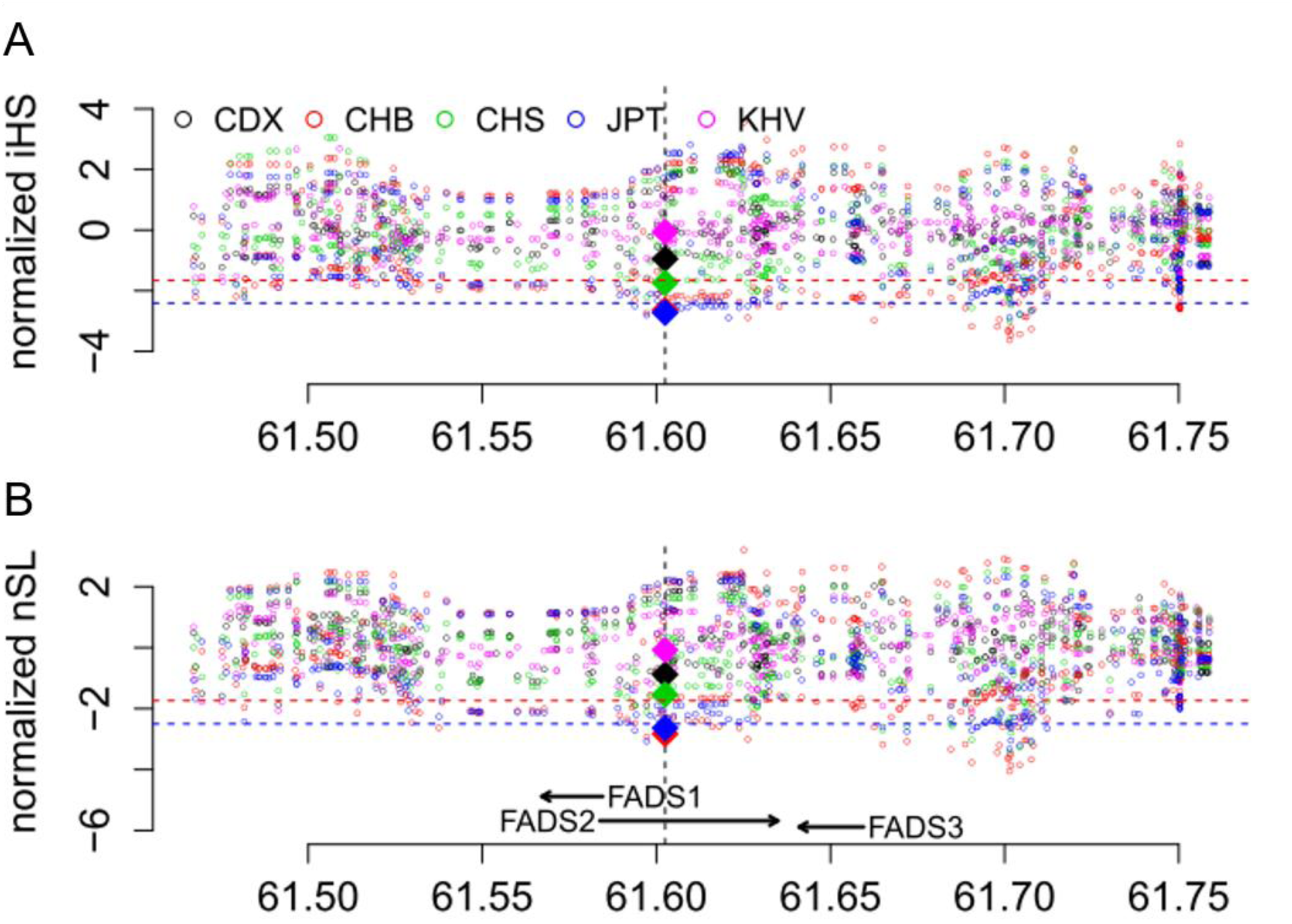
Patterns of normalized iHS and nSL across a 300 Kb region around FADS genes in five East Asian populations. The two statistics were calculated for each of the common SNPs (minor allele frequency > 5%) in the region. The values for rs66698963 in each population are highlighted as diamonds filled with population-specific colors. The red dashed lines represent empirical *p* values of 0.05, while the blue dashed lines represent empirical *p* values of 0.01 based on estimates of all genome-wide common SNPs. These cutoffs are almost the same across all five populations.

**Supplementary Figure S18.**
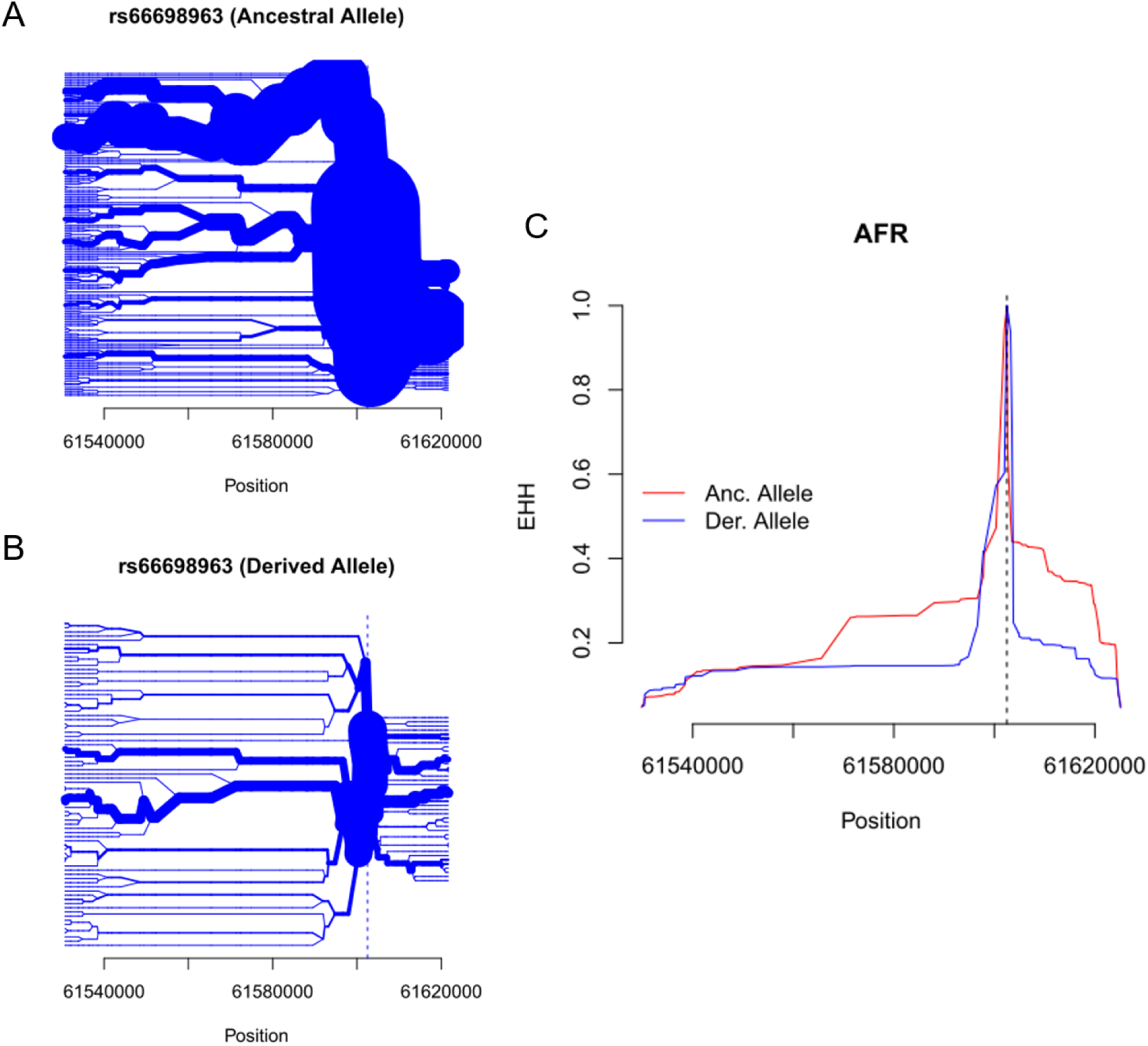
The Insertion allele of rs66698963 is associated with signals of positive selection in populations of African origin. A) Haplotype bifurcation diagrams for the Insertion allele at rs66698963; B) Haplotype bifurcation diagrams for the Deletion allele at rs66698963; C) The plot of expanded haplotype homozygosity (EHH) for rs66698963. The EHH values are plotted against the physical distance extending both upstream and downstream of the target SNP. The ancestral allele refers to the Insertion and the derived allele refers to the Deletion allele.

**Supplementary Figure S19.**
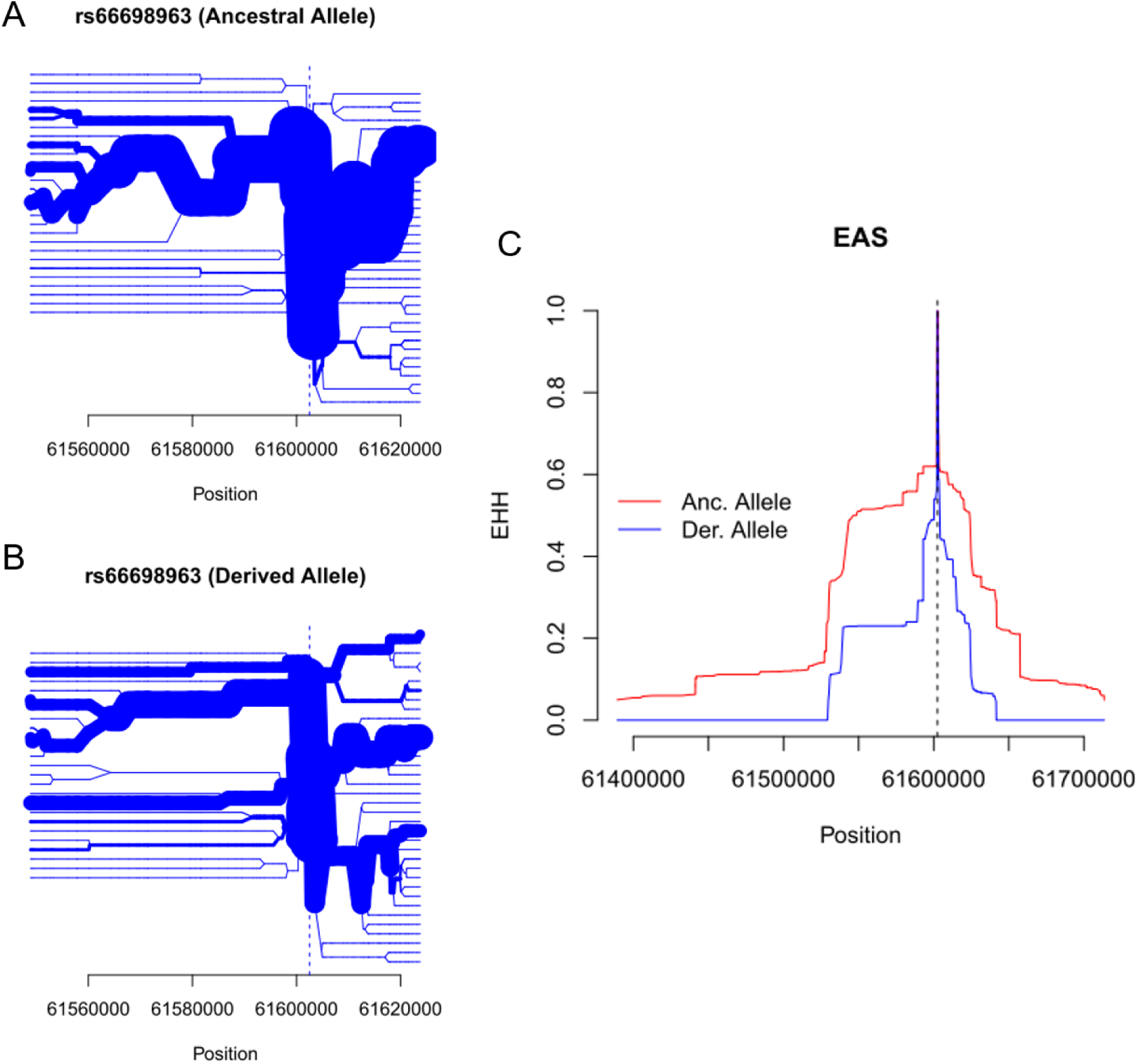
The Insertion allele of rs66698963 is associated with signals of positive selection in populations of East Asian origin. A) Haplotype bifurcation diagrams for the Insertion allele at rs66698963; B) Haplotype bifurcation diagrams for the Deletion allele at rs66698963; C) The plot of expanded haplotype homozygosity (EHH) for rs66698963. The EHH values are plotted against the physical distance extending both upstream and downstream of the target SNP. The ancestral allele refers to the Insertion and the derived allele refers to the Deletion allele.

**Supplementary Figure S20.**
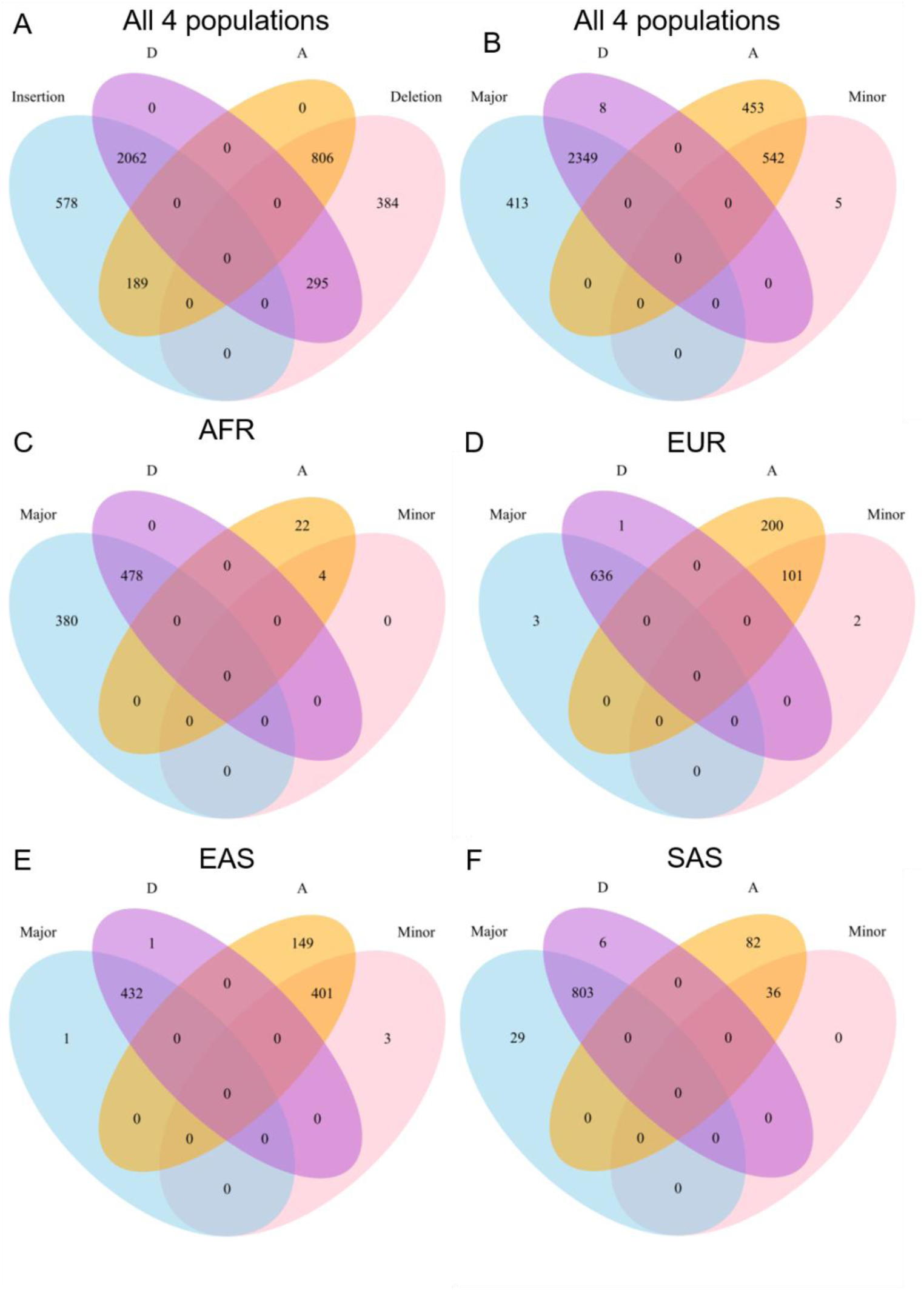
Strong associations between D haplotype and Insertion allele (Major), and between A haplotype and Deletion allele (Minor). A) Venn diagrams representing overlaps of chromosomes carrying D haplotype, A haplotype, Insertion allele and Deletion allele in all four continental populations. Venn diagrams representing overlaps of chromosomes carrying D haplotype, A haplotype, Major haplotype and Minor haplotype in B) all four continental populations; C) Africans; D) Europeans; E) East Asians; F) South Asians. D and A haplotypes are defined with 28 SNPs while Major and Minor haplotypes are defined with 10 SNPs. They have three SNPs in common. Venn diagrams are not proportional to the real numbers represented.

**Supplementary Figure S21.**
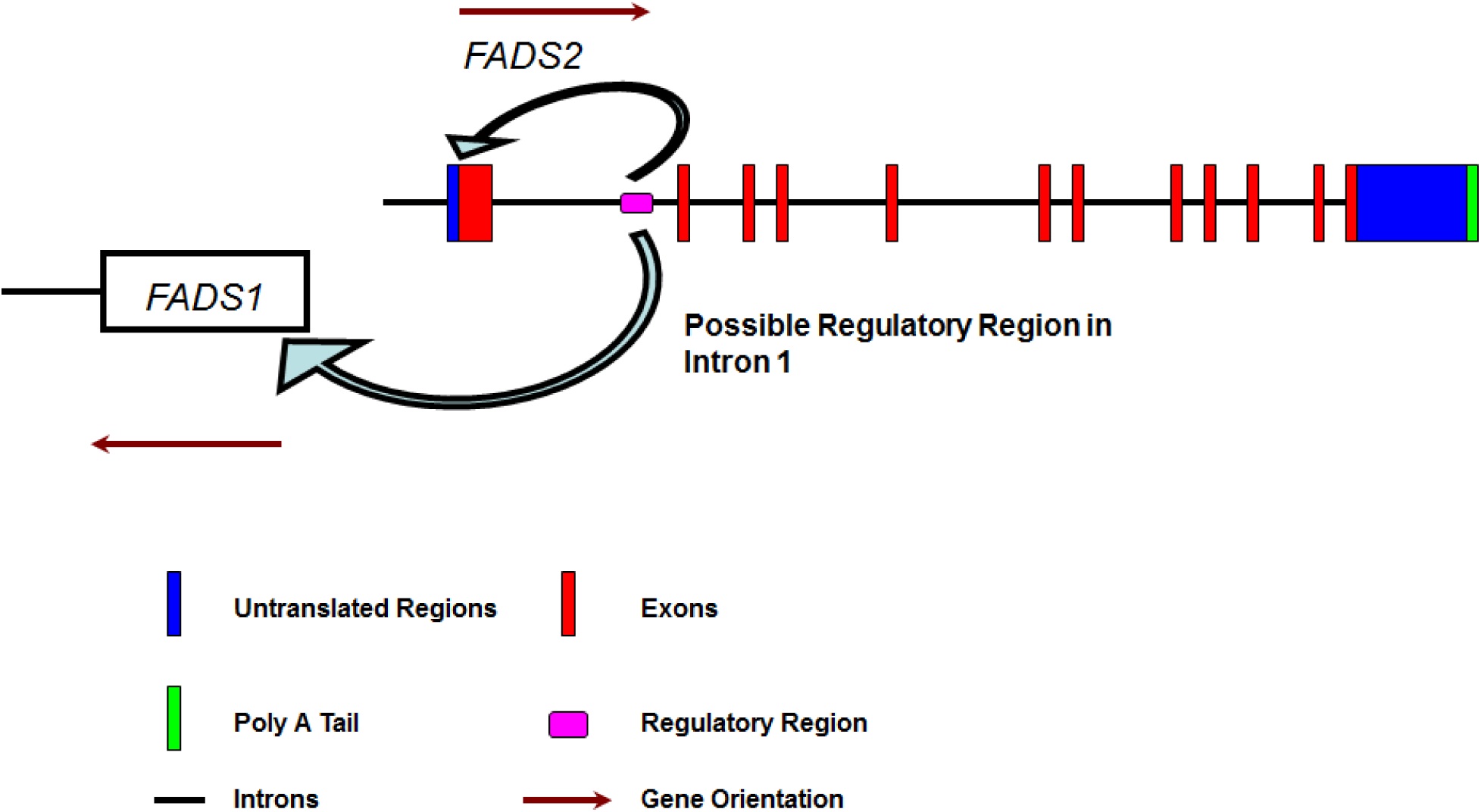
Cartoon illustration of the conserved genomic region in intron 1 of FADS2. The regulatory region includes rs66698963 near the SRE that regulates expression of FADS1 and FADS2.

**Supplementary Figure S22.**
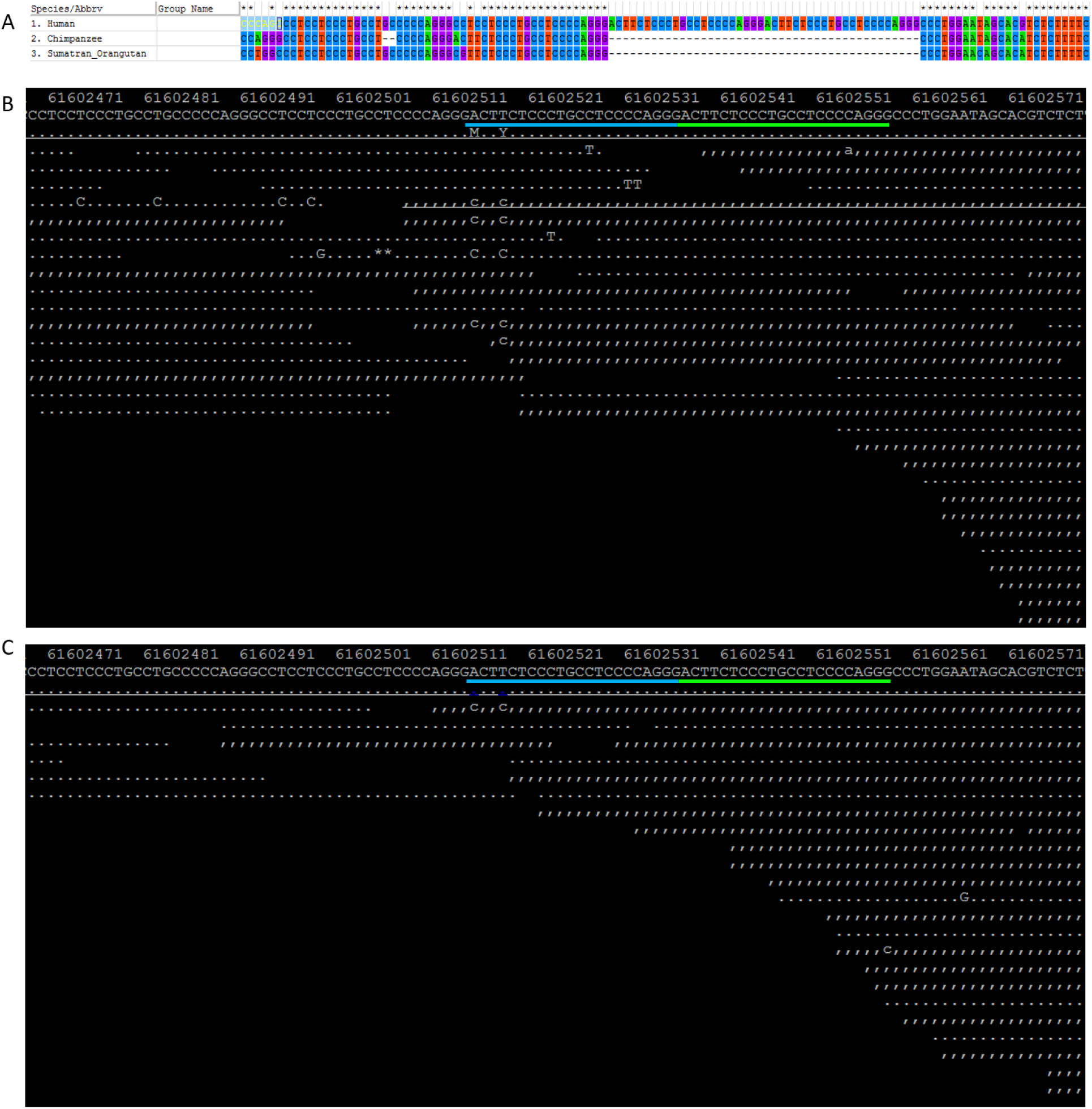
The ancestral allele of rs66698963. A) Alignment results surrounding rs66698963 for human, Chimpanzee and Orangutan. Neither the insertion allele, nor the deletion allele of rs66698963, is present in Chimpanzee and Orangutan; B) Mapping results of sequencing reads from a Neanderthal from the Altai Mountains to the Modern Human genome. This Neanderthal individual carries the insertion allele; C) Mapping results of sequencing reads from an archaic Denisovan individual to the Modern Human genome. The Denisovan individual carries the insertion allele.

## References

Alvheim AR, Malde MK, Osei-Hyiaman D, Lin YH, Pawlosky RJ, Madsen L, Kristiansen K, Froyland L, Hibbeln JR. 2012. Dietary linoleic acid elevates endogenous 2-AG and anandamide and induces obesity. Obesity (Silver Spring) 20:1984–1994.

Ameur A, Enroth S, Johansson A, Zaboli G, Igl W, Johansson AC, Rivas MA, Daly MJ, Schmitz G, Hicks AA, et al. 2012. Genetic adaptation of fatty-acid metabolism: a human-specific haplotype increasing the biosynthesis of long-chain omega-3 and omega-6 fatty acids. Am J Hum Genet 90:809–820.

Aulchenko YS, Ripatti S, Lindqvist I, Boomsma D, Heid IM, Pramstaller PP, Penninx BW, Janssens AC, Wilson JF, Spector T, et al. 2009. Loci influencing lipid levels and coronary heart disease risk in 16 European population cohorts. Nat Genet 41:47–55.

Auton A, Brooks LD, Durbin RM, Garrison EP, Kang HM, Korbel JO, Marchini JL, McCarthy S, McVean GA, Abecasis GR. 2015. A global reference for human genetic variation. Nature 526:68–74.

Blasbalg TL, Hibbeln JR, Ramsden CE, Majchrzak SF, Rawlings RR. 2011. Changes in consumption of omega-3 and omega-6 fatty acids in the United States during the 20th century. Am J Clin Nutr 93:950–962.

Bokor S, Dumont J, Spinneker A, Gonzalez-Gross M, Nova E, Widhalm K, Moschonis G, Stehle P, Amouyel P, De Henauw S, et al. 2010. Single nucleotide polymorphisms in the FADS gene cluster are associated with delta-5 and delta-6 desaturase activities estimated by serum fatty acid ratios. J Lipid Res 51:2325–2333.

Brenna JT, Lapillonne A. 2009. Background paper on fat and fatty acid requirements during pregnancy and lactation. Ann Nutr Metab 55:97–122.

Cao C, Pressman EK, Cooper EM, Guillet R, Westerman M, O’Brien KO. 2014. Placental heme receptor LRP1 correlates with the heme exporter FLVCR1 and neonatal iron status. Reproduction 148:295–302.

Carlson SE, Colombo J, Gajewski BJ, Gustafson KM, Mundy D, Yeast J, Georgieff MK, Markley LA, Kerling EH, Shaddy DJ. 2013. DHA supplementation and pregnancy outcomes. Am J Clin Nutr 97:808–815.

Caspi A, Williams B, Kim-Cohen J, Craig IW, Milne BJ, Poulton R, Schalkwyk LC, Taylor A, Werts H, Moffitt TE. 2007. Moderation of breastfeeding effects on the IQ by genetic variation in fatty acid metabolism. Proc Natl Acad Sci U S A 104:18860–18865.

Diau GY, Hsieh AT, Sarkadi-Nagy EA, Wijendran V, Nathanielsz PW, Brenna JT. 2005. The influence of long chain polyunsaturate supplementation on docosahexaenoic acid and arachidonic acid in baboon neonate central nervous system. BMC Med 3:11.

Eaton SB, Eaton SB, 3rd, Sinclair AJ, Cordain L, Mann NJ. 1998. Dietary intake of long-chain polyunsaturated fatty acids during the paleolithic. World Rev Nutr Diet 83:12–23.

Emken EA, Adlof RO, Gulley RM. 1994. Dietary linoleic acid influences desaturation and acylation of deuterium-labeled linoleic and linolenic acids in young adult males. Biochim Biophys Acta 1213:277–288.

Fay JC, Wu CI. 2000. Hitchhiking under positive Darwinian selection. Genetics 155:1405–1413.

Ferrer-Admetlla A, Liang M, Korneliussen T, Nielsen R. 2014. On detecting incomplete soft or hard selective sweeps using haplotype structure. Mol Biol Evol 31:1275–1291.

Ferretti A, Nelson GJ, Schmidt PC, Kelley DS, Bartolini G, Flanagan VP. 1997. Increased dietary arachidonic acid enhances the synthesis of vasoactive eicosanoids in humans. Lipids 32:435–439.

Folch J, Lees M, Sloane Stanley GH. 1957. A simple method for the isolation and purification of total lipides from animal tissues. J Biol Chem 226:497–509.

Fumagalli M, Moltke I, Grarup N, Racimo F, Bjerregaard P, Jorgensen ME, Korneliussen TS, Gerbault P, Skotte L, Linneberg A, et al. 2015. Greenlandic Inuit show genetic signatures of diet and climate adaptation. Science 349:1343–1347.

Gadgil M, Joshi K, Pandit A, Otiv S, Joshi R, Brenna JT, Patwardhan B. 2014. Imbalance of folic acid and vitamin B12 is associated with birth outcome: an Indian pregnant women study. Eur J Clin Nutr 68:726–729.

Gautier M, Vitalis R. 2012. rehh: an R package to detect footprints of selection in genome-wide SNP data from haplotype structure. Bioinformatics 28:1176–1177.

Gibson RA, Neumann MA, Lien EL, Boyd KA, Tu WC. 2013. Docosahexaenoic acid synthesis from alpha-linolenic acid is inhibited by diets high in polyunsaturated fatty acids. Prostaglandins Leukot Essent Fatty Acids 88:139–146.

Gibson RA, Sinclair AJ. 1981. Are Eskimos obligate carnivores? Lancet 1:1100.

Groen-Blokhuis MM, Franic S, van Beijsterveldt CE, de Geus E, Bartels M, Davies GE, Ehli EA, Xiao X, Scheet PA, Althoff R, et al. 2013. A prospective study of the effects of breastfeeding and FADS2 polymorphisms on cognition and hyperactivity/attention problems. Am J Med Genet B Neuropsychiatr Genet 162B:457–465.

Hester AG, Murphy RC, Uhlson CJ, Ivester P, Lee TC, Sergeant S, Miller LR, Howard TD, Mathias RA, Chilton FH. 2014. Relationship between a common variant in the fatty acid desaturase (FADS) cluster and eicosanoid generation in humans. J Biol Chem 289:22482–22489.

Illig T, Gieger C, Zhai G, Romisch-Margl W, Wang-Sattler R, Prehn C, Altmaier E, Kastenmuller G, Kato BS, Mewes HW, et al. 2010. A genome-wide perspective of genetic variation in human metabolism. Nat Genet 42:137–141.

Key TJ, Appleby PN, Rosell MS. 2006. Health effects of vegetarian and vegan diets. Proc Nutr Soc 65:35–41.

Lattka E, Rzehak P, Szabo E, Jakobik V, Weck M, Weyermann M, Grallert H, Rothenbacher D, Heinrich J, Brenner H, et al. 2011. Genetic variants in the FADS gene cluster are associated with arachidonic acid concentrations of human breast milk at 1.5 and 6 mo postpartum and influence the course of milk dodecanoic, tetracosenoic, and trans-9-octadecenoic acid concentrations over the duration of lactation. Am J Clin Nutr 93:382–391.

Li H, Handsaker B, Wysoker A, Fennell T, Ruan J, Homer N, Marth G, Abecasis G, Durbin R. 2009. The Sequence Alignment/Map format and SAMtools. Bioinformatics 25:2078–2079.

Li Z, Wu X, He B, Zhang L. 2014. Vindel: a simple pipeline for checking indel redundancy. BMC Bioinformatics 15:359.

Liu B, Chen J, Shen B. 2011. Genome-wide analysis of the transcription factor binding preference of human bi-directional promoters and functional annotation of related gene pairs. BMC Syst Biol 5 Suppl 1:S2.

Lu JT, Wang Y, Gibbs RA, Yu F. 2012. Characterizing linkage disequilibrium and evaluating imputation power of human genomic insertion-deletion polymorphisms. Genome Biol 13:R15.

Malerba G, Schaeffer L, Xumerle L, Klopp N, Trabetti E, Biscuola M, Cavallari U, Galavotti R, Martinelli N, Guarini P, et al. 2008. SNPs of the FADS gene cluster are associated with polyunsaturated fatty acids in a cohort of patients with cardiovascular disease. Lipids 43:289–299.

Marquardt A, Stohr H, White K, Weber BH. 2000. cDNA cloning, genomic structure, and chromosomal localization of three members of the human fatty acid desaturase family. Genomics 66:175–183.

Martinelli N, Girelli D, Malerba G, Guarini P, Illig T, Trabetti E, Sandri M, Friso S, Pizzolo F, Schaeffer L, et al. 2008. FADS genotypes and desaturase activity estimated by the ratio of arachidonic acid to linoleic acid are associated with inflammation and coronary artery disease. Am J Clin Nutr 88:941–949.

Mathias RA, Fu W, Akey JM, Ainsworth HC, Torgerson DG, Ruczinski I, Sergeant S, Barnes KC, Chilton FH. 2012. Adaptive evolution of the FADS gene cluster within Africa. PLoS One 7:e44926.

Meyer M, Kircher M, Gansauge MT, Li H, Racimo F, Mallick S, Schraiber JG, Jay F, Prufer K, de Filippo C, et al. 2012. A high-coverage genome sequence from an archaic Denisovan individual. Science 338:222–226.

Morrison WR, Smith LM. 1964. Preparation of Fatty Acid Methyl Esters and Dimethylacetals from Lipids with Boron Fluoride–Methanol. J Lipid Res 5:600–608.

Naqvi AZ, Davis RB, Mukamal KJ. 2012. Dietary fatty acids and peripheral artery disease in adults. Atherosclerosis 222:545–550.

Nelson GJ, Kelley DS, Emken EA, Phinney SD, Kyle D, Ferretti A. 1997. A human dietary arachidonic acid supplementation study conducted in a metabolic research unit: rationale and design. Lipids 32:415–420.

Nussbaum RL, McInnes RR, Willard HF, Hamosh A, Thompson MW. 2007. Thompson & Thompson genetics in medicine. Philadelphia: Saunders/Elsevier.

O’Dea K, Sinclair AJ. 1985. The effects of low-fat diets rich in arachidonic acid on the composition of plasma fatty acids and bleeding time in Australian aborigines. J Nutr Sci Vitaminol (Tokyo) 31:441–453.

Park HG, Kothapalli KS, Park WJ, DeAllie C, Liu L, Liang A, Lawrence P, Brenna JT. 2016. Palmitic acid (16:0) competes with omega-6 linoleic and omega-3 a-linolenic acids for FADS2 mediated Delta6-desaturation. Biochim Biophys Acta 1861:91–97.

Park HG, Park WJ, Kothapalli KS, Brenna JT. 2015. The fatty acid desaturase 2 (FADS2) gene product catalyzes Delta4 desaturation to yield n-3 docosahexaenoic acid and n-6 docosapentaenoic acid in human cells. FASEB J 29:3911–3919.

Park WJ, Kothapalli KS, Lawrence P, Brenna JT. 2011. FADS2 function loss at the cancer hotspot 11q13 locus diverts lipid signaling precursor synthesis to unusual eicosanoid fatty acids. PLoS One 6:e28186.

Park WJ, Kothapalli KS, Lawrence P, Tyburczy C, Brenna JT. 2009. An alternate pathway to long-chain polyunsaturates: the FADS2 gene product Delta8-desaturates 20:2n-6 and 20:3n-3. J Lipid Res 50:1195–1202.

Park WJ, Kothapalli KS, Reardon HT, Kim LY, Brenna JT. 2009. Novel fatty acid desaturase 3 (FADS3) transcripts generated by alternative splicing. Gene 446:28–34.

Park WJ, Kothapalli KS, Reardon HT, Lawrence P, Qian SB, Brenna JT. 2012. A novel FADS1 isoform potentiates FADS2-mediated production of eicosanoid precursor fatty acids. J Lipid Res 53:1502–1512.

Park WJ, Reardon HT, Tyburczy C, Kothapalli KS, Brenna JT. 2010. Alternative splicing generates a novel FADS2 alternative transcript in baboons. Mol Biol Rep 37:2403–2406.

Pollner R, Schmidt C, Fischer G, Kuhn K, Poschl E. 1997. Cooperative and competitive interactions of regulatory elements are involved in the control of divergent transcription of human Col4A1 and Col4A2 genes. FEBS Lett 405:31–36.

Prufer K, Racimo F, Patterson N, Jay F, Sankararaman S, Sawyer S, Heinze A, Renaud G, Sudmant PH, de Filippo C, et al. 2014. The complete genome sequence of a Neanderthal from the Altai Mountains. Nature 505:43–49.

Raheja BS, Sadikot SM, Phatak RB, Rao MB. 1993. Significance of the N-6/N-3 ratio for insulin action in diabetes. Ann N Y Acad Sci 683:258–271.

Reardon HT, Zhang J, Kothapalli KS, Kim AJ, Park WJ, Brenna JT. 2012. Insertion-deletions in a FADS2 intron 1 conserved regulatory locus control expression of fatty acid desaturases 1 and 2 and modulate response to simvastatin. Prostaglandins Leukot Essent Fatty Acids 87:25–33.

Remaley AT, Bark S, Walts AD, Freeman L, Shulenin S, Annilo T, Elgin E, Rhodes HE, Joyce C, Dean M, et al. 2002. Comparative genome analysis of potential regulatory elements in the ABCG5-ABCG8 gene cluster. Biochem Biophys Res Commun 295:276–282.

Rzehak P, Thijs C, Standl M, Mommers M, Glaser C, Jansen E, Klopp N, Koppelman GH, Singmann P, Postma DS, et al. 2010. Variants of the FADS1 FADS2 gene cluster, blood levels of polyunsaturated fatty acids and eczema in children within the first 2 years of life. PLoS One 5:e13261.

Sabeti PC, Varilly P, Fry B, Lohmueller J, Hostetter E, Cotsapas C, Xie X, Byrne EH, McCarroll SA, Gaudet R, et al. 2007. Genome-wide detection and characterization of positive selection in human populations. Nature 449:913–918.

Schaeffer L, Gohlke H, Muller M, Heid IM, Palmer LJ, Kompauer I, Demmelmair H, Illig T, Koletzko B, Heinrich J. 2006. Common genetic variants of the FADS1 FADS2 gene cluster and their reconstructed haplotypes are associated with the fatty acid composition in phospholipids. Hum Mol Genet 15:1745–1756.

Sergeant S, Hugenschmidt CE, Rudock ME, Ziegler JT, Ivester P, Ainsworth HC, Vaidya D, Case LD, Langefeld CD, Freedman BI, et al. 2012. Differences in arachidonic acid levels and fatty acid desaturase (FADS) gene variants in African Americans and European Americans with diabetes or the metabolic syndrome. Br J Nutr 107:547–555.

Sharma S, Zhou X, Thibault DM, Himes BE, Liu A, Szefler SJ, Strunk R, Castro M, Hansel NN, Diette GB, et al. 2014. A genome-wide survey of CD4(+) lymphocyte regulatory genetic variants identifies novel asthma genes. J Allergy Clin Immunol 134:1153–1162.

Simopoulos AP. 2002. The importance of the ratio of omega-6/omega-3 essential fatty acids. Biomed Pharmacother 56:365–379.

Smuts CM, Huang M, Mundy D, Plasse T, Major S, Carlson SE. 2003. A randomized trial of docosahexaenoic acid supplementation during the third trimester of pregnancy. Obstet Gynecol 101:469–479.

Standl M, Sausenthaler S, Lattka E, Koletzko S, Bauer CP, Wichmann HE, von Berg A, Berdel D, Kramer U, Schaaf B, et al. 2011. FADS gene variants modulate the effect of dietary fatty acid intake on allergic diseases in children. Clin Exp Allergy 41:1757–1766.

Steer CD, Hibbeln JR, Golding J, Davey Smith G. 2012. Polyunsaturated fatty acid levels in blood during pregnancy, at birth and at 7 years: their associations with two common FADS2 polymorphisms. Hum Mol Genet 21:1504–1512.

Szpiech ZA, Hernandez RD. 2014. selscan: an efficient multithreaded program to perform EHH-based scans for positive selection. Mol Biol Evol 31:2824–2827.

Tajima F. 1989. Statistical method for testing the neutral mutation hypothesis by DNA polymorphism. Genetics 123:585–595.

Tanaka T, Shen J, Abecasis GR, Kisialiou A, Ordovas JM, Guralnik JM, Singleton A, Bandinelli S, Cherubini A, Arnett D, et al. 2009. Genome-wide association study of plasma polyunsaturated fatty acids in the InCHIANTI Study. PLoS Genet 5:e1000338.

Voight BF, Kudaravalli S, Wen X, Pritchard JK. 2006. A map of recent positive selection in the human genome. PLoS Biol 4:e72.

Weir BS, Cockerham CC. 1984. Estimating F-Statistics for the Analysis of Population Structure. Evolution 38:1358–1370.

Wood KE, Lau A, Mantzioris E, Gibson RA, Ramsden CE, Muhlhausler BS. 2014. A low omega-6 polyunsaturated fatty acid (n-6 PUFA) diet increases omega-3 (n-3) long chain PUFA status in plasma phospholipids in humans. Prostaglandins Leukot Essent Fatty Acids 90:133–138.

Xie L, Innis SM. 2008. Genetic variants of the FADS1 FADS2 gene cluster are associated with altered (n-6) and (n-3) essential fatty acids in plasma and erythrocyte phospholipids in women during pregnancy and in breast milk during lactation. J Nutr 138:2222–2228.

Yi X, Liang Y, Huerta-Sanchez E, Jin X, Cuo ZX, Pool JE, Xu X, Jiang H, Vinckenbosch N, Korneliussen TS, et al. 2010. Sequencing of 50 human exomes reveals adaptation to high altitude. Science 329:75–78.

Zail SS, Pickering A. 1979. Fatty acid composition of erythrocytes in hereditary spherocytosis. Br J Haematol 42:399–402.

Zhang B, Jia WH, Matsuda K, Kweon SS, Matsuo K, Xiang YB, Shin A, Jee SH, Kim DH, Cai Q, et al. 2014. Large-scale genetic study in East Asians identifies six new loci associated with colorectal cancer risk. Nat Genet 46:533–542.

Zhang X, Johnson AD, Hendricks AE, Hwang SJ, Tanriverdi K, Ganesh SK, Smith NL, Peyser PA, Freedman JE, O’Donnell CJ. 2014. Genetic associations with expression for genes implicated in GWAS studies for atherosclerotic cardiovascular disease and blood phenotypes. Hum Mol Genet 23:782–795.

